# Ca^2+^ and Membrane Potential Transitions During Action Potentials are Self-Similar to Each Other and to Variability of AP Firing Intervals Across the Broad Physiologic Range of AP Intervals During Autonomic Receptor Stimulation

**DOI:** 10.1101/2020.09.01.277756

**Authors:** Dongmei Yang, Christopher H. Morrell, Alexey E. Lyashkov, Syevda Tagirova, Ihor Zahanich, Yael Yaniv, Tatiana M. Vinogradova, Bruce D. Ziman, Victor A. Maltsev, Edward G. Lakatta

**Affiliations:** Laboratory of Cardiovascular Science, National Institute on Aging, National Institutes of Health, Baltimore, USA; Mathematics and Statistics Department, Loyola University Maryland, Baltimore, USA; Biomedical Engineering Faculty, Technion-IIT, Haifa, Israel

**Keywords:** single sinoatrial nodal pacemaker cells, Local diastolic Ca^2+^ releases, diastolic depolarization, autonomic receptor stimulation, self-similarity of Ca^2+^ and membrane potential during action potentials, action potential, firing interval variability

## Abstract

Ca^2+^ and V_m_ transitions occurring throughout AP cycles in sinoatrial nodal (SAN) cells are **cues** that: (1) **not only regulate** activation states of molecules operating within criticality (Ca^2+^ domain) and limit-cycle (V_m_ domain) mechanisms of a coupled-clock system that underlies SAN cell automaticity; (2) but are also **regulated by** the activation states of the clock molecules they regulate. In other terms, these cues are **both** causes and effects of clock molecular activation (recursion). Recently, we demonstrated that Ca^2+^ and V_m_ transitions during AP cycles in single SAN cells isolated from mice, guinea pigs, rabbits and humans are self-similar (obey a power law) and are also self-similar to trans-species AP firing intervals of these cells in *vitro*, to heart rate in *vivo*, and to body mass.

Neurotransmitter stimulation of β adrenergic receptor or cholinergic receptor initiated signaling in SAN cells modulates their AP firing rate and rhythm by impacting on the degree to which SAN clocks couple to each other, creating the broad physiologic range of SAN cell mean AP firing intervals and firing interval variabilities. Here we show that Ca^2+^ and V_m_ domain kinetic transitions (time to AP ignition in diastole and 90% AP recovery) occurring within given AP, the mean AP firing intervals, and AP firing interval variabilities within time-series of APs in 230 individual SAN cells are self-similar (obey power laws). In other terms, these long-range correlations inform on self-similar distributions of order among SAN cells across the entire broad physiologic range of SAN AP firing intervals, regardless of whether autonomic receptors of these cells are stimulated or not, and regardless of the type (adrenergic or cholinergic) of autonomic receptor stimulation. These long-range correlations among distributions of Ca^2+^ and V_m_ kinetic functions that regulate SAN cell clock coupling during each AP cycle in different **individual**, isolated SAN cells not in contact with each other. Our numerical model simulations further extended our perspectives to the molecular scale and demonstrated that many ion currents also behave self-similar across autonomic states.

Thus, to ensure rapid flexibility of AP firing rates in response to different types and degrees of autonomic input, nature “did not reinvent molecular wheels within the coupled-clock system of pacemaker cells”, but differentially engaged or scaled the kinetics of gears that regulate the rate and rhythm at which the “wheels spin” in a given autonomic input context.

## Introduction

The heart is a central player within a hierarchical system of clocks operating within the autonomic neuro-visceral axis that creates and synchronizes rhythmic functions ranging from msec to days and beyond (Lakatta, 2021; Shivkumar et al., 2016). The heart’s beating rate and rhythm are regulated by autonomic input to sinoatrial nodal (SAN) pacemaker cells that modulates functions within a coupled-clock system intrinsic to SAN cells (Lakatta, Maltsev, & Vinogradova, 2010).

### What is the coupled-clock system within pacemaker cells and how do clocks couple to each other?

The SAN cell coupled-clock system is comprised of a calcium “clock”, the sarcoplasmic reticulum (SR), that continuously oscillates Ca^2+^ via a *criticality* mechanism (Nivala, Ko, Nivala, Weiss, & Qu, 2012) and phase-like transitions (A.V. Maltsev et al., 2011); the Ca^2+^ clock is continuously but variably coupled to a “membrane clock”, an ensemble of surface membrane ion channels that generates current oscillations via a *limit-cycle* mechanism (Weiss & Qu, 2020). The criticality mechanisms, in turn, are governed by power law and self-similarity across wide scales (Bak, 1999). The “biochemical engine” of the coupled-clock system is a constitutively active, Ca^2+^ calmodulin-dependent adenylyl cyclase (AC) that generates cyclic AMP, leading to modulation of cAMP-gated ion channels, EPAC signaling, and PKA and CAMKII-dependent kinase activities, mechanisms that regulate intracellular Ca^2+^ levels, Ca^2+^ dynamics and membrane potential within SAN cells (Lakatta, Maltsev, Bogdanov, Stern, & Vinogradova, 2003; Lakatta et al., 2010; Lakatta et al., 2006; Lakatta, Vinogradova, & Maltsev, 2008; V. A. Maltsev & Lakatta, 2008; Yaniv et al., 2015). Variable rates and rhythms at which SAN cells fire APs are controlled by the kinetics of sub-cellular and cell-wide transitions in [Ca^2+^] gradients and the membrane potential (V_m_), and the extent to which V_m_ and Ca^2+^ become coupled during AP cycles in any given epoch. For more details, see Supplementary Discussion.

The well-known variability of AP firing intervals of isolated SAN cells in *vitro*, of SAN tissue *ex vivo* or of heartbeat intervals in *vivo* (Monfredi et al., 2014; Yaniv, Ahmet, et al., 2014) indicates that coupled-clock system Ca^2+^ and V_m_ functions during AP cycles never achieve a true steady state from one AP to the next.

These time-dependent Ca^2+^ and V_m_ domain transitions during APs are **cues**, that **not only regulate** activation states of clock molecules, but are also **regulated by** the activation status of the very molecules they regulate. In other terms, changes in these cues cause changes in clock molecule activation that feedback to change the characteristics of activation cues. This recursive dynamic imparts robustness to SAN cell automaticity (Lyashkov, Behar, Lakatta, Yaniv, & Maltsev, 2018; V. A. Maltsev & Lakatta, 2009). The variability in the degree to which Ca^2+^ and membrane clock molecules couple to each other throughout AP cycles is due to transitions (changes) that occur in Ca^2+^ and V_m_ domain cues throughout AP cycles (Monfredi et al., 2013; Yaniv, Lyashkov, et al., 2014).

Spontaneous transitions in sub-cellular Ca^2+^ and V_m_ domains that emerge during the spontaneous diastolic depolarization (DD) phase of an AP cycle have been conceptualized as the AP “ignition phase” (Lyashkov et al., 2018). The ignition process in the Ca^2+^ domain is linked to the emergence of local spontaneous, diastolic oscillatory RyR activation, that generates local Ca^2+^ releases (LCRs) that self-organize to form Ca^2+^ wavelets that propagate locally (Bogdanov, Vinogradova, & Lakatta, 2001; Vinogradova et al., 2004). Ca^2+^-dependent activation of the surface membrane electrogenic Na^+^/Ca^2+^ exchanger generates inward current that accelerates diastolic V_m_ depolarization and couples the clocks. The time at which the rate of this feed-forward crosstalk acutely accelerates to 0.15V/s, marks the onset of the coupled-clock ignition process (Lyashkov et al., 2018).

Following ignition onset, the extent to which the Ca^2+^ and V_m_ clock become coupled continues to increase throughout the diastolic period as LCRs and Ca^2+^ wavelets emerge at remote areas across the cell and continue to self-organize in time throughout the cellular space, creating an explosive ensemble Ca^2+^ signal that progressively depolarizes the cell membrane, i.e., clock-coupling progressively increases. This Ca^2+^-induced change in V_m_ increase in clock-coupling cues the activation of low-voltage activated Ca^2+^ channels (Ca_v_1.3 and Ca_v_3.1), resulting in Ca^2+^ influx that contributes to the further organization of the ensemble LCR Ca^2+^ signal via feed-forward electrochemical (Ca^2+^-V_m_-Ca^2+^) signaling, when the diastolic V_m_ enters a range that cues the activation of L type Ca^2+^ channels (Ca_v_1.2). The ignition phase of the coupled-Ca^2+^ and V_m_ domain sub-cellular kinetic transitions culminates in the generation of cell-wide events; a marked transition in the rate of V_m_ depolarization, due to the activation of Ca_v_1.2, results in the rapid AP upstroke and Ca^2+^ influx, which, via Ca^2+^-induced Ca^2+^ release from the SR via RyRs, generates an AP-induced cytosolic Ca^2+^ transient. In other terms, spontaneous, cell-wide Ca^2+^ signals and APs in SAN cells emerge from spatio-temporal self-organization of spontaneous sub-cellular Ca^2+^ oscillations (the *criticality* mechanism) (Nivala et al., 2012). Serca 2a pumping Ca^2+^ into SR and K^+^ channel repolarization of V_m_ return the Ca^2+^ and V_m_ domain cues toward their diastolic levels at which LCRs again begin to emerge, creating the ignition phase of the next AP cycle.

### Self-organized criticality

Spatiotemporal self-organization across geometric scales (sub-cellular to cell-wide) is a manifestation of criticality that has been observed in excitable cells throughout nature (Stožer et al., 2019) including cultured astrocytes (Jung, Cornell-Bell, Madden, & Moss, 1998), immature oocytes (Lopez, Piegari, Sigaut, & Dawson, 2012) and mouse cardiac ventricular myocytes (Nivala et al., 2012). Self-similar, scale-free distributions of parameters across wide scales that obey power law behavior (are ln-ln linear) are an indication of their self-ordered criticality (Bak, 1999).

It has recently been discovered that coupling of sub-cellular Ca^2+^ signals (cues) generated by the Ca^2+^ clock within isolated SAN cells to the cell surface membrane proteins during APs to elicit a change in V_m_ manifests long range power law correlations (are self-similar) across species (S. T. Sirenko et al., 2021). Specifically, Ca^2+^ and V_m_ domain kinetic transitions (cues) during AP cycles in single SAN cells isolated from mice, guinea pigs, rabbits and humans are self-similar to each other during APS and self-similar to trans-species AP firing intervals of these cells in *vitro*, to heart rate in *vivo*, and to body mass (S. T. Sirenko et al., 2021).

Neurotransmitter stimulation of βAR or CR-initiated signaling modulates SAN cells’ AP firing rate and rhythm by impacting on coupled-clock protein functions, modulating the degree to which criticality (Ca^2+^ domain) and limit-cycle (V_m_ domain) mechanisms couple to each other during AP cycles (Lakatta et al., 2010; V. A. Maltsev & Lakatta, 2009). AP firing intervals in rabbit SAN cells during autonomic stimulation vary over a four-fold range, from about 200 msec during βAR stimulation (βARs) up to about 800 msec during CR stimulation (CRs) (Lyashkov et al., 2009; Vinogradova, Bogdanov, & Lakatta, 2002).

We hypothesized that transitions in V_m_ and Ca^2+^ domain cues during the diastolic AP ignition (Lyashkov et al., 2018) and recovery phases (Figure 1) of APs are: (1) not only self-similar to each other in cells without autonomic receptor stimulation (control cells), but are self-similar to V_m_ and Ca^2+^ cues in **other** cells during CRs and during βARs; and (2) that Ca^2+^ and V_m_ cues during APs are self-similar to AP firing interval variabilities (and therefore self-similar to mean AP firing intervals) regardless of the presence or absence or type of autonomic receptor stimulation. In other terms, we hypothesized that Ca^2+^ and V_m_ domain clock-coupling cues occurring during **all** APs are self-similar to each other i.e., manifest long-range correlations **in all** isolated SAN cells within populations of cells that differ with respect to autonomic input and, that these Ca^2+^ and V_m_ cues during APs are also self-similar to the rate and rhythm of AP firing across the **entire range** of AP firing intervals created by these cues in **all** isolated SAN cells.

**Figure 1.**
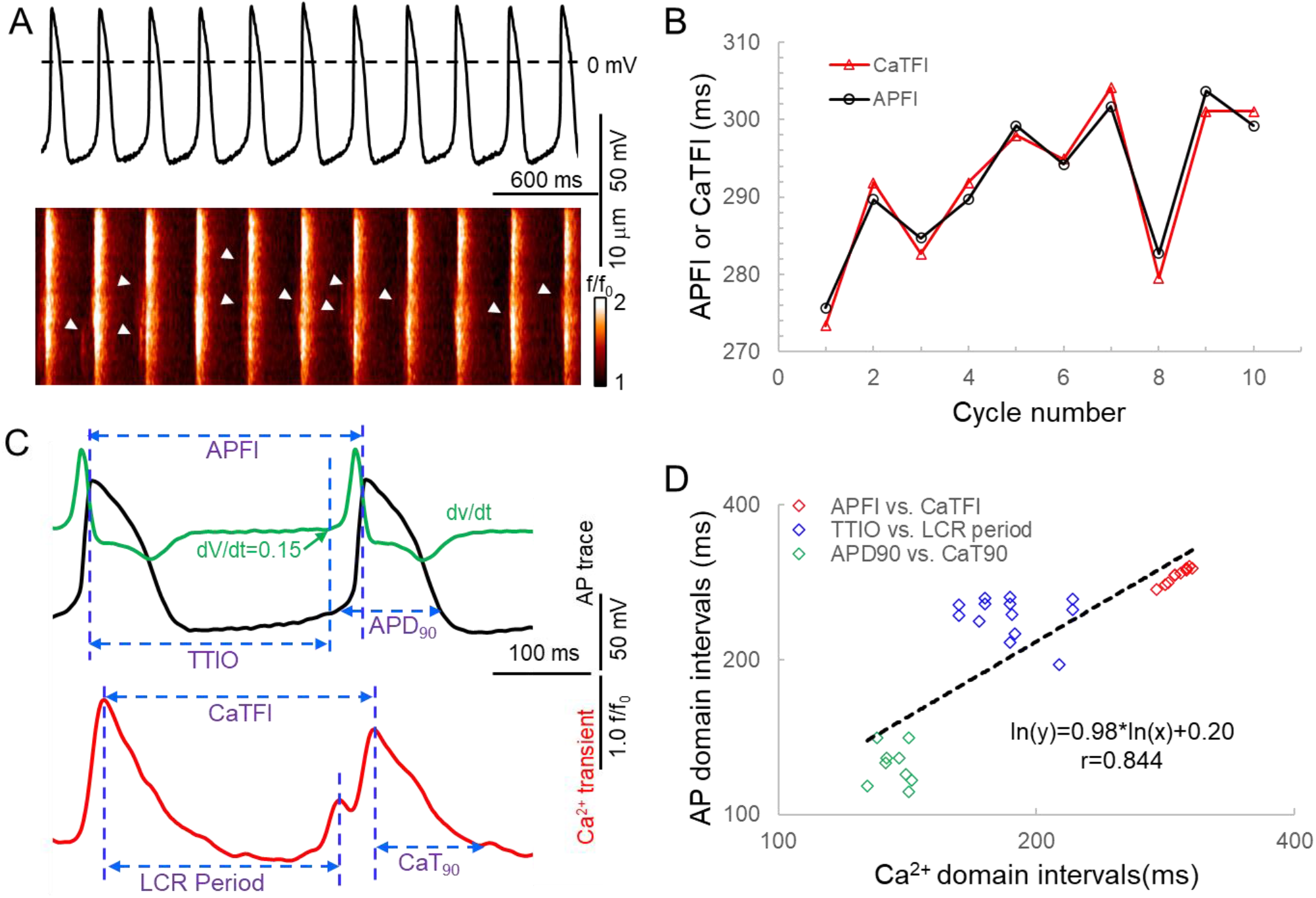
Simultaneous recording of AP and Ca^2+^ in a 10 beat time series of APs (A) and the measured APFI and CaTFI (B); the definition of parameters measured in V_m_ and Ca^2+^ domains (C), and the self-similarity of the Vm and Ca^2+^ parameters during APs to each other and to the APFI (D).

To test these hypotheses: we studied a large population (n=230) of single rabbit SAN cells to which we applied: CRs (carbachol, CCh), to one subset of cells; βARs (isoproterenol, ISO) to another subset; and no autonomic receptor stimulation to a third subset of cells. This created three populations of SAN cells having APFIs distributed across the entire physiologic range. We measured intracellular Ca^2+^ or membrane potential in these cells to: (1) characterize the times to ignition onset, and times to 90% recovery of V_m_ and Ca^2+^ parameters during APs in AP time-series; and (2), to determine the correlations of these V_m_ and Ca^2+^ kinetic parameters to each other during APs, to AP firing interval variability (and therefore to mean AP firing intervals). Thus, the data set to be analyzed consisted of 12 different kinetic parameters in each cell population (control, CCh and ISO); 6 parameter means, 3 each in the Ca^2+^ and V_m_ domains; and 6 parameter variabilities (SDs) around the means. To determine the degree of self-similarity among V_m_ and Ca^2+^ domain parameters, we constructed density distribution plots and applied correlation, power law, and principal component (PC) analyses to Ca^2+^ and V_m_ domain data sets separately, and to the combined Ca^2+^ and V_m_ data sets. We further extended our perspectives from cell population and single cell levels downwards to the molecular scale by performing numerical modeling simulation and analyzing variabilities of ion currents and Ca^2+^ with respect to APFI to determine whether these ion currents and Ca^2+^ also obeyed a power law across autonomic states.

## Materials and Methods

The study was performed in accordance with the Guide for the Care and Use of Laboratory Animals published by the National Institutes of Health (NIH Publication number. 85-23, revised 1996). The experimental protocols have been approved by the Animal Care and Use Committee of the National Institutes of Health (protocol #034LCS2016). Materials and methods briefly presented here are detailed in Supplementary Material.

### Single, isolated rabbit SAN cells isolation

Single, spindle-shaped, spontaneously beating SAN cells were isolated from the hearts of New Zealand rabbits (Charles River Laboratories, Wilmington, MA) as described previously (Vinogradova TM, 2000).

### Spontaneous APs recordings

Time series of spontaneous APs were recorded in subsets of freshly isolated SAN cells using the perforated patch-clamp technique with Axopatch 200B patch-clamp amplifier (Axon Instruments) (Bogdanov et al., 2001) at 34+0.5°C. AP parameters (Figure 1) were measured via a customized program (Lyashkov et al., 2018) were APFI, APD_90_, and the time to ignition onset (TTIO) measured by the time at which diastolic membrane potential dV/dt accelerates to 0.15 V/sec (Figure 1) which reflects the onset of the ignition phase of the AP cycle (Lyashkov et al., 2018).

### Ca^2+^ measurements

In another subset of SAN cells, AP-induced global Ca^2+^ transients and spontaneous LCRs (Figure 1) were measured at 34+0.5°C with a confocal microscope (Zeiss LSM510, Germany) in the line-scan mode (Vinogradova et al., 2004; D. Yang, Lyashkov, Li, Ziman, & Lakatta, 2012). The interval between the peaks of two adjacent AP-induced Ca^2+^ transients (Figure 1) is defined as Ca^2+^ transient (CaT) firing interval (CaTFI), which is highly correlated with the APFI, as demonstrated by simultaneous recordings of V_m_ and Ca^2+^ in a separate subset of cells (Figure 1). The LCR period is defined as the time from the peak of the prior AP-induced Ca^2+^ transient to an LCR peak in diastole (Figure 1); the time to 90% decay of the CaT was defined as CaT_90_.

### Numerical modeling

We performed numerical simulations using a modified Maltsev-Lakatta model that features the coupled-clock mechanism (V. A. Maltsev & Lakatta, 2009). The computer code for the original model is freely available and can be downloaded and run in CellML format (http://models.cellml.org/workspace/maltsev_2009) using the Cellular Open Resource software developed by Alan Garny at Oxford University in the UK (Garny, Noble, Hunter, & Kohl, 2009) (for recent development of this software see http://www.opencor.ws/). The original model could not be directly used for APFI variability simulations because it is a system of first-order differential equations that is deterministic and showing no APFI variability in limit cycle oscillatory regime of AP steady firing. Thus, we modified the model to generate variability of AP waveforms by supplementing total membrane current (I_tot_) with an additional randomly fluctuating current around its zero-mean value, known as perturbation current or I_per_ (as previously implemented by Henggui Zhang (Monfredi et al., 2014)). Furthermore, we also performed an additional set of simulations with I_per_ added to Ca^2+^ release flux to mimic the effect of stochastic LCRs (Bogdanov et al., 2006; Monfredi et al., 2013). Using the resultant stochastic dynamical system of SAN cell, we simulated fluctuating APFI, ion currents, and Ca^2+^ dynamics for three conditions: (i) basal AP firing, (ii) during βARs with ISO (100 nM), and (iii) during CRs with CCh (100 nM). The effects of autonomic modulation (conditions ii and iii) were modelled as previously described (V. A. Maltsev & Lakatta, 2010), except modulation of I_CaL_ current by CCh that was modelled as described by Zaza et al. (Zaza, Robinson, & Difrancesco, 1996). All model equations and parameters are provided in Supplementary Material.

### Experimental design and statistics

Supplementary Figure S1 illustrates schematic of the experimental design to assess long-range correlations of V_m_ and Ca^2+^ parameters during APs and APFI intervals in cells within and among populations of cells that differed with respect to autonomic input. V_m_ and Ca^2+^ parameter intervals (msec) are presented as Mean ± SD. AP firing interval variability within a time series is taken as standard deviation (SD) about the mean, or as the coefficient of variation (CV, the ratio of SD to the mean).

Analyses of V_m_ and Ca^2+^ parameter interval distributions measured in AP time-series in **different** cells determined the association among pairs of variables using Spearman’s correlations for average data, and Pearson’s correlation for both average and individual data (Howell, 2002); In cells in which Ca^2+^ was measured, the mean interval between AP- induced CaTs was usually longer than that the mean APFI in cells in which APs were recorded (due to slight buffering effects of the fluorescent Ca^2+^ probe). To allow all the variables to be combined into a single analysis, we matched on the APFI variable Z-scores in control, ISO, or CCh populations by ‘matchit’ function in R (Ho, Imai, King, & Stuart, 2011).

Density estimates of **within cell** standard deviations and means of parameters of cells within each autonomic state population, are presented as nonparametric kernel estimates of probability density functions, scaled so that the total area under each curve is unity (Silverman, 1986).

In order to determine whether distributions of parameter means and SDs are self-similar, i.e., obeyed a power law suggesting fractal-like behavior, we constructed ln-ln plots of distributions of the AP and Ca^2+^ function means and SDs measured across the broad range of apparent steady states in the absence of, or the presence of βARs or CRs. (Kucera, Heuschkel, Renaud, & Rohr, 2000; Yaniv, Lyashkov, & Lakatta, 2013).

The relationships among all distributions of all parameters (means and SDs), were also assessed in principal component analyses (Johnson & Wichern, 2008).

In a few cells in which experimentally measured parameter interval distributions measured in the V_m_ and Ca^2+^ domains in the same cell and for numerical simulation of ion currents and Ca^2+^ prior to and during autonomic receptor stimulation Poincaré indices were employed to define long-range correlations among variables. A Poincaré Plot graphs a parameter (*n*), in an AP time series on the *x*-axis versus the same parameter of the succeeding AP (n+1) on the *y*-axis, i.e. one takes a sequence of parameters and plots each one against the following parameter (Huikuri et al., 2000). When statistical inference was performed, a p-value<0.05 was considered statistically significant.

## Results

### Assessment of self-similarity of Ca^2+^ and V_m_ kinetic interval parameters to each other during APs and to AP firing interval variability

Figure 1 A illustrates a time-series of APs in a SAN cell during which Ca^2+^ and V_m_ were simultaneously measured in the absence of autonomic receptor stimulation. Kinetic transitions in Ca^2+^ and V_m_ parameters as illustrated in Figure 1B, were assessed during each AP. Figure 1C shows that V_m_ and Ca^2+^ parameters during AP time series are self-similar to each other and are also self-similar to the AP firing intervals (r=0.844 in this cell).

Figure 2 illustrates the time series of AP intervals in Figure 1A plotted as phase-plane diagrams in which V_m_ is depicted as a function of Ca^2+^ throughout each cycle. The times at which various channels are activated throughout the V_m_/Ca loop are indicated. The times of ignition and 90% recovery are also indicated. The point resolution is 3.072ms, and the point spread indicates the rates at which the electrochemical signal changes during each cycle. The dashed line in the figure marks the border between the disordered and ordered molecular activation. The arrows indicate the direction of the electrochemical signal emergence.

**Figure 2.**
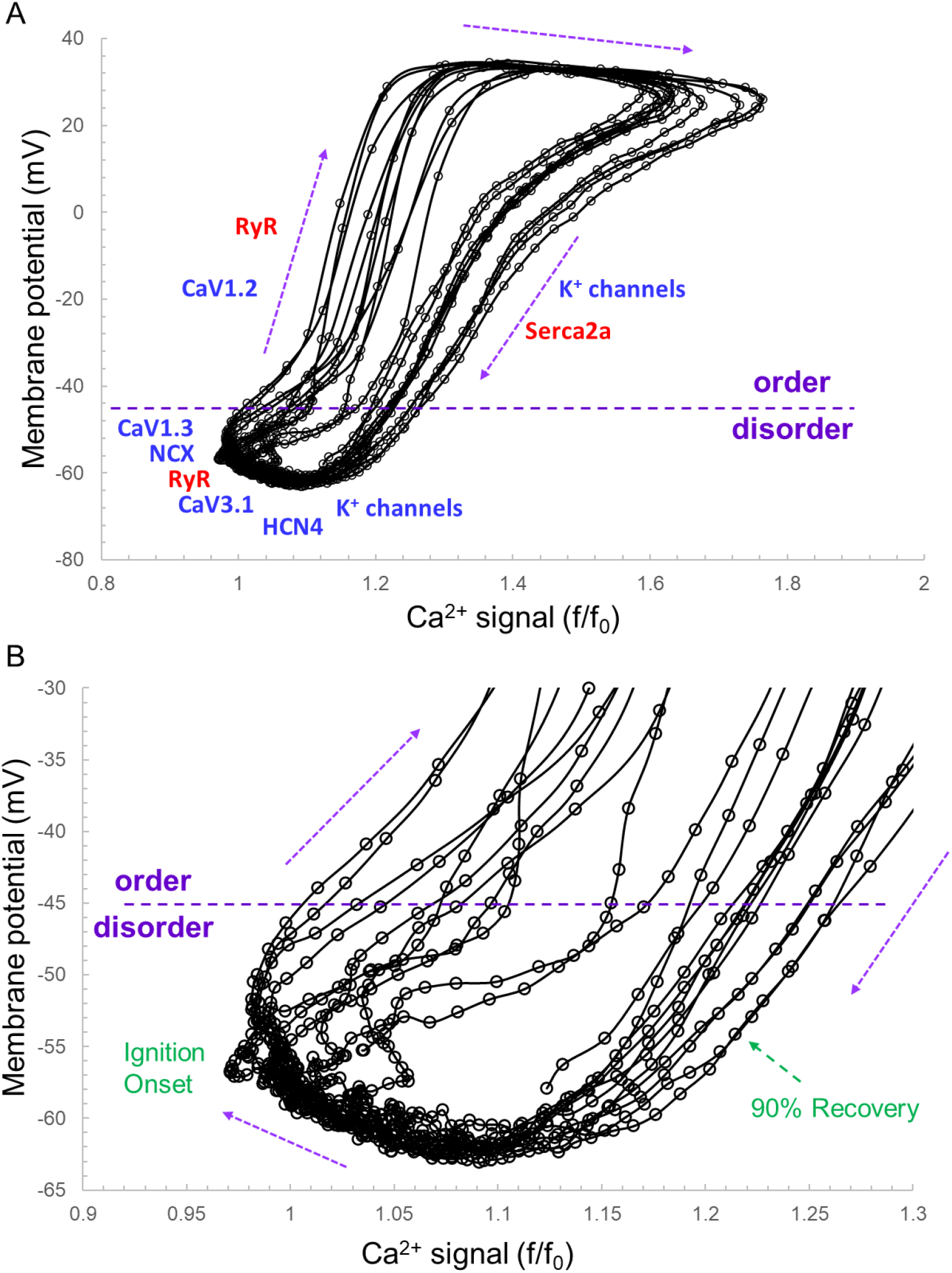
A illustrates the time series of AP intervals in Figure 1A plotted as phase-plane diagrams in which V_m_ is depicted as a function of Ca^2+^ throughout each cycle. The times at which various channels are activated throughout the V_m_/Ca phase-plane loop are indicated. The times of ignition and 90% recovery are also indicated (B). The point resolution is 3.072 ms, and the spread indicates the rates at which the electrochemical signal changes during each cycle. The dashed line in the figure marks the border between the disordered and ordered molecular activation. The arrows indicate the direction of the electrochemical signal emergence.

A Poincaré plot (scatter graph) constructed from consecutive data points in a time-series (Figure 3A) is a convenient tool that provides information on correlations (self-similarity) of data across the time-series. The X axis defines the parameter (n) occurrence in msec, and the Y axis defines the parameter occurrence at (n+1). The Poincaré plot in Figure 3A depicts the data of the time-series of the cell in Figure 1. Note that although the means vary over a three-fold range, all six parameter means (the Ca^2+^ and V_m_ interval parameters measured during APs and AP firing intervals) in the absence of autonomic receptor stimulation are described by a line of identity, indicating their self-similarity across the AP time series. Quantitative analysis of short and long term variability in a given time series of observations entails fitting an ellipse to each cloud of data points within the Poincaré Plot (Figure 3B): The length of a line describing the slope of the long axis of each ellipse is referred to as SD2 of the data points (c.f. Figure inset); the length of the line describing the slope of the short axis, which is perpendicular in direction to the long axis line, is referred to as SD1. Note in Figure 3C that the SD1 is self-similar to SD2 across the four-fold range of Ca^2+^ and V_m_ parameters. The center point of each ellipse i.e., the intersection of SD1 and SD2, is the **average** interval between events (AP intervals or other parameters measured in the time series) within the time series. SD1/SD2 (Figure inset) informs on non-linear trends (unequal lengths of SD1 and SD2) across intervals within each ellipse.

**Figure 3.**
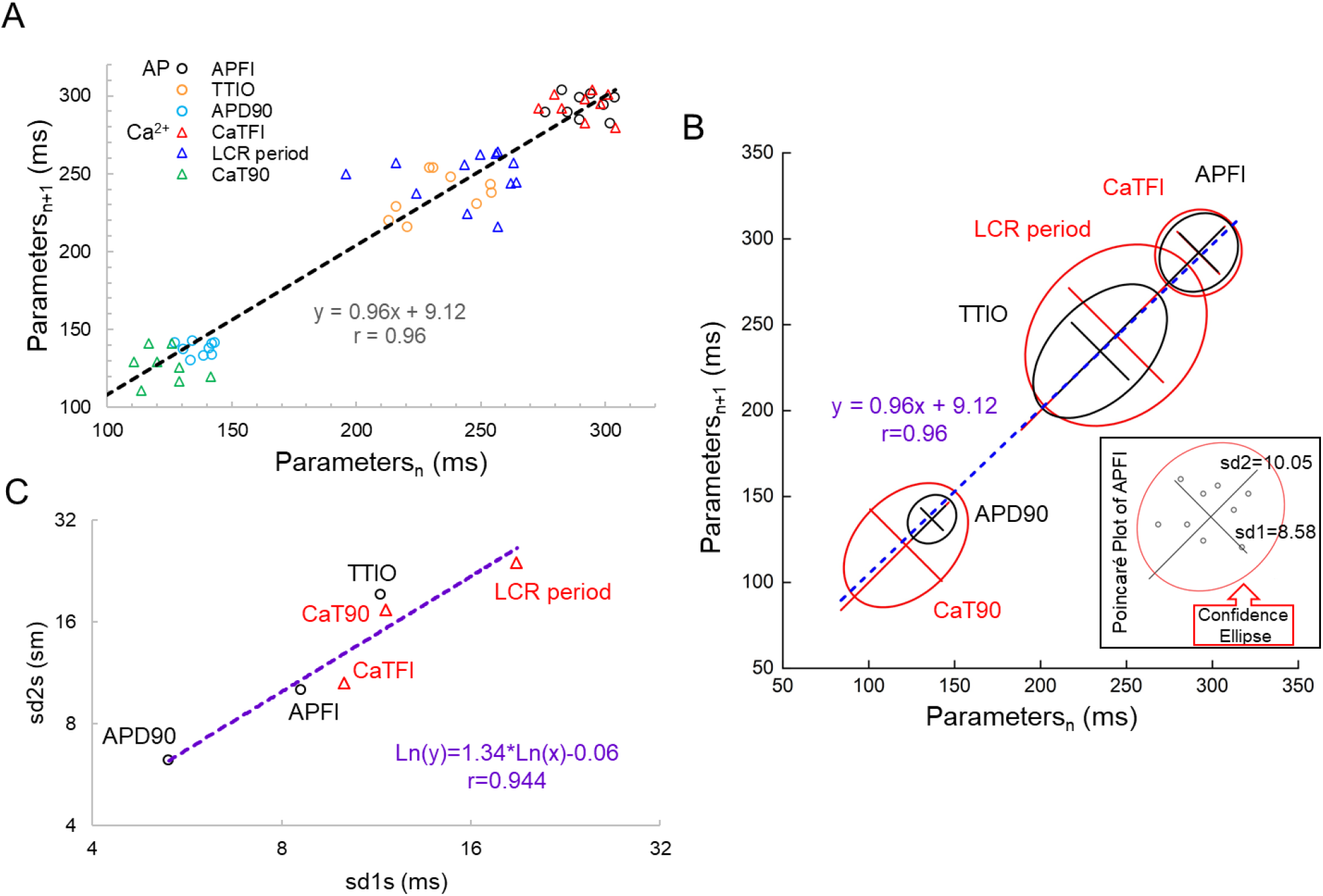
Poincare plots (A) and fitting of the Poincare plot Ellipse clouds (B), the relation of sd2s to sd1s (C) of the 6 parameters from simultaneously recorded during the 10 beats time series from Figure 1.

Figure 4 illustrates combined Poincaré plots of TTIO, APD_90_, during APs and AP firing intervals in time-series of APs of two representative cells: one cell in control and during CRs by CCh; and the other cell in control and during βARs stimulation by ISO.

**Figure 4.**
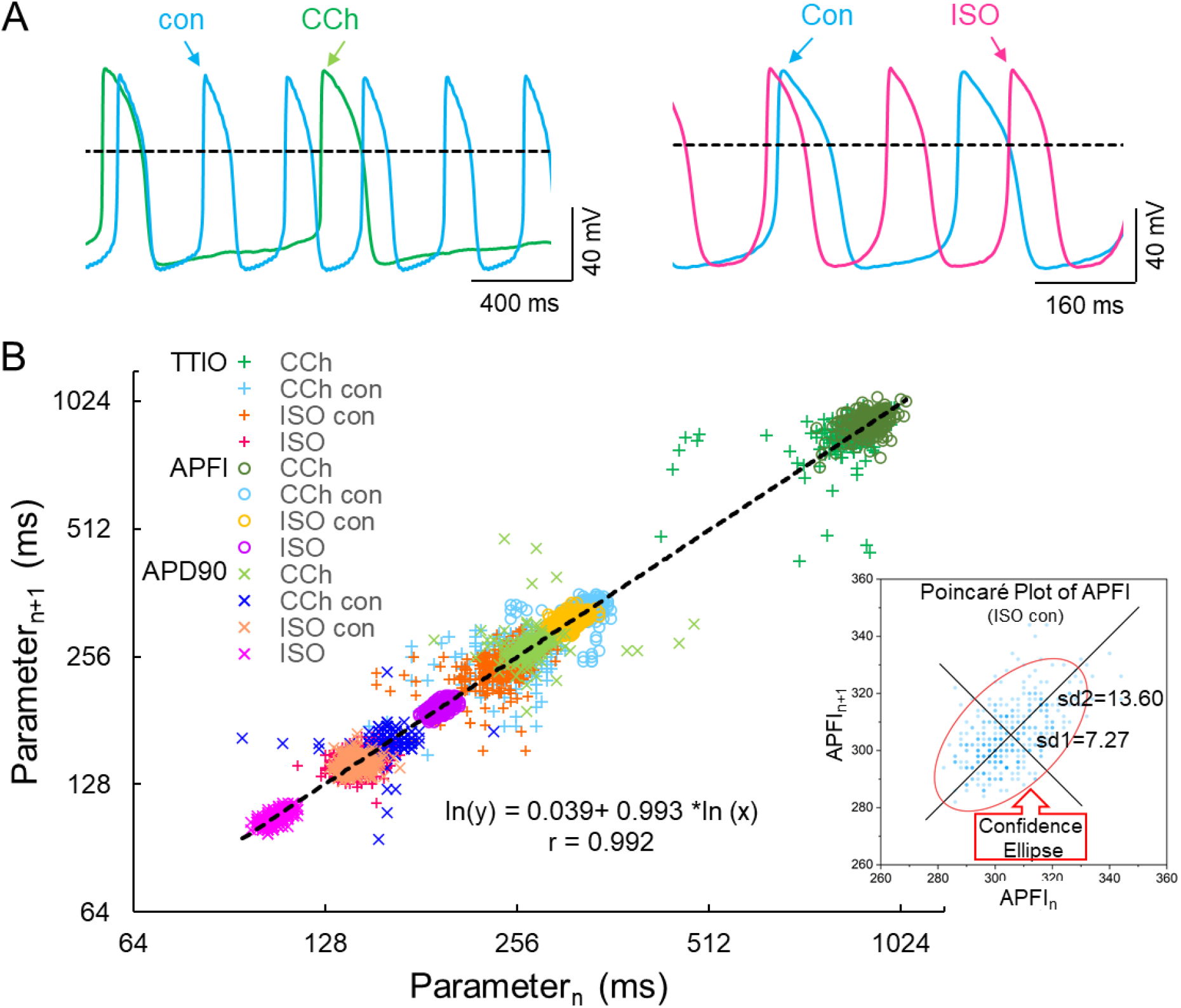
Self-control AP recordings in 2 cells during CCh or ISO (A), Poincare plots of the 3 measured V_m_ domain parameters in 3 autonomic states: CCh, control (2 cells) and ISO (B, number of beats in each time series was 197, 397, 397 and 900, respectively, in carbochol, carbochol control, isoproterenol control and isoproterenol). The inset shows an example of Ellipse fitting.

Although the range of absolute values of kinetic interval parameters of cells depicted in the Poincaré Plot in Figure 4B vary by 20-fold, all points (n=5673) are self-similar, i.e. are fit by a single line (r=0.992) with a slope of unity, passing nearly through the origin.

Supplementary Table S1 lists the SD1s, SD2s, SD1/SD2, the means of the TTIO and APD_90_ intervals and AP firing intervals depicted in Figure 4B. Note also that the SD1s, SD2s, and SD1/SD2 of TTIO, APD_90_ and AP firing intervals progressively increase from ISO to control, and **markedly** increase from control to CCh, creating degrees of non-linearity across the combined control, ISO and CCh states, which is also reflected in the mean AP firing intervals across the three states (Supplementary Table S1). The V_m_ transitions during APs across different autonomic states are self-similar to APFIs across these states (Figure 5A). Figure 5B shows the self-similarity of parameter means of SD1s of V_m_ parameters to their SD2s across autonomic states, indicating self-similarity of short term (e.g., beat to beat) and long term (e.g., rhythm across more than 2 beats) variabilities across autonomic states within the time series. Figure 5C shows the self-similarity of all parameter means depicted in Figures 4, 5A and 5B to their SDs across autonomic states.

**Figure 5.**
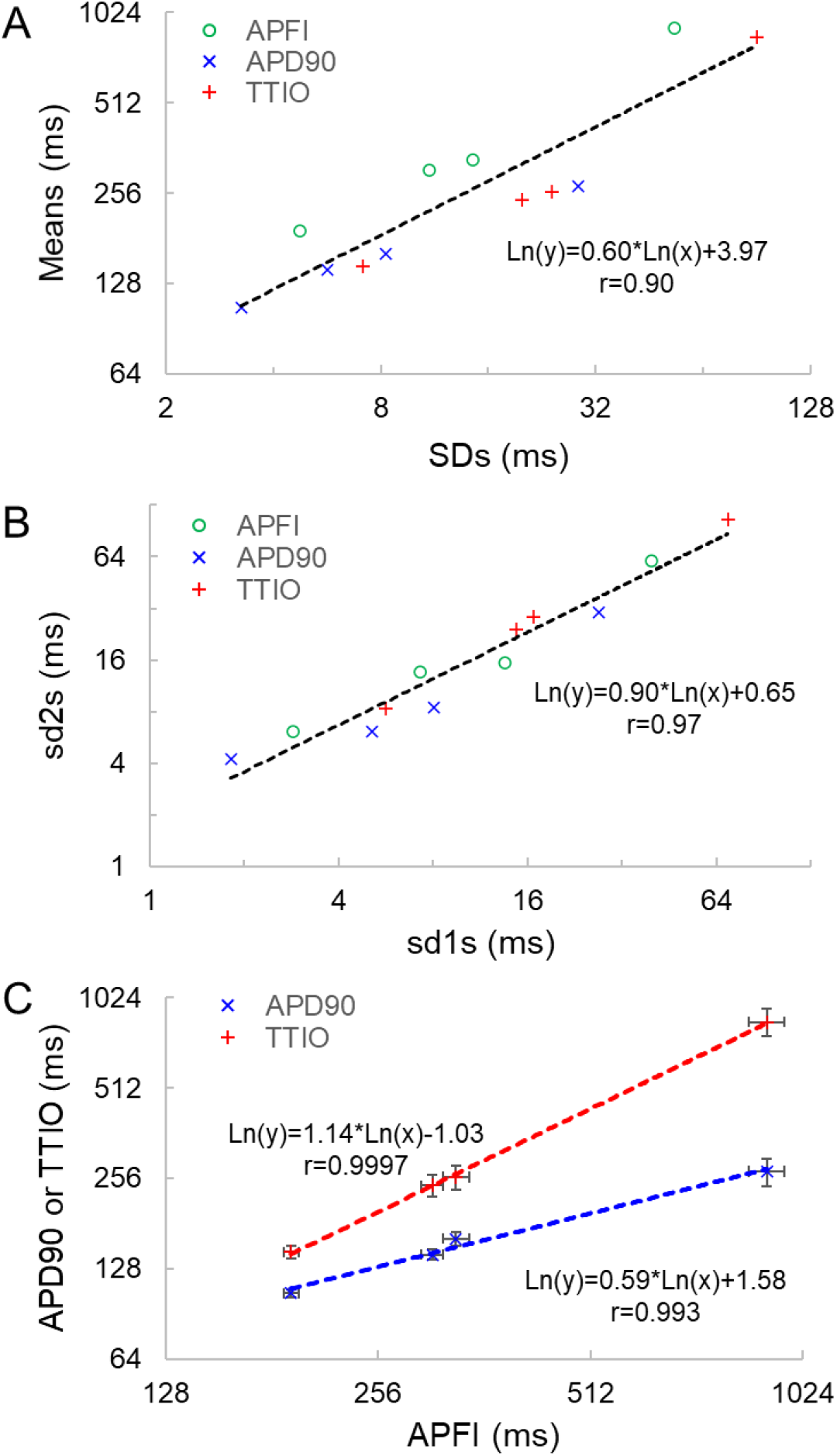
Correlation between Mean and SD of the 3 Vm parameters (A), correlation of Poincare Ellipse fitting sd1 and sd2 (B), and the self-similarity of APD90 and TTIO to APFI (C) across the 3 autonomic states, the same as in figure 4.

The data in Figures 4 and 5 demonstrate that over the entire range of physiologic AP firing intervals from 192.7ms in ISO to 305.5 ms in control, and to 910.3 ms in CCh, variabilities of TTIO and APD_90_ measured <u>during</u> APs are self-similar to each other and are also self-similar to the variability of AP firing intervals within the time-series, and therefore self-similar to the mean AP firing interval of the time-series.

Self-similarity among V_m_ variables in the cells in Figure 4 across autonomic states in control and during ISO and CCh in Figures 2 and 3A can also easily be ascertained from the shapes of their population density distributions (Figure 6). Note that in the cell super-fused with CCh the distributions of TTIO, APD_90_ and AP firing intervals (Panels A-C) are broader than in control or during ISO. Note also that the distributions of kinetic transitions during APs and APFI become more synchronized from CCh to ISO (Panels A-C). In other terms, the degree to which V_m_ and Ca^2+^ parameters are synchronized during APs increases from CCh to control to ISO, similar to the AP firing variabilities and mean AP firing intervals.

**Figure 6.**
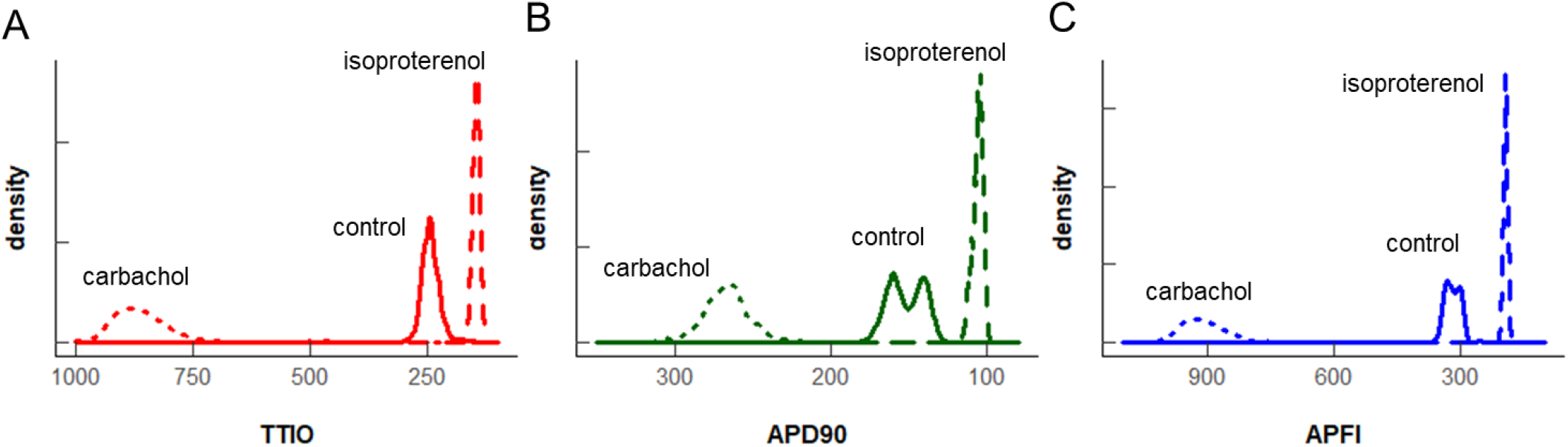
Density distributions of the self-control Vm parameters measured in the same cell as in figure 4 and 5 (2 control cells are combined). The density distributions are presented as nonparametric kernel estimates of probability density functions (Silverman, 1986), scaled so that the total area within each curve is unity.

### Self-Similarity of Ca^2+^ and V_m_ parameter <u>means</u> to each other during APs and to APFI variabilities and mean AP firing intervals across autonomic states in <u>different</u> cells

We next determined whether the self-similarity (long-range correlations) of Ca^2+^ to V_m_ parameters measured within the same cells as depicted in Figures 1–6 extends to populations of **different** cells within and among different autonomic states. To accomplish this, we applied CRs (carbachol, CCh), to one subset of cells; βAR stimulation (isoproterenol, ISO) to another subset; and no autonomic receptor stimulation to a third subset of cells. This created populations of SAN cells having APFIs distributed across the entire physiologic range.

Table 1 lists descriptive statistics (means and SDs) of the variabilities and means of Ca^2+^ and V_m_ kinetic parameter intervals measured in **different** cells within and among populations of cells that **differ** with respect to autonomic input. The **mean SD** of V_m_ and Ca^2+^ domain parameters listed in Table 1A gives the average time-series variability of each parameter among cells within **each** of the three cell populations (control, ISO, or CCh). The standard deviation of the SDs (**SDSD**) in Table 1A tells us how variable the SDs of each parameter are among cells within **each** cell population. The **means of each parameter** measured within a time series (Table 1B) tells us the **average level** of the parameter among cells within each of the three populations; and the **SD of the means** tells us the **variability of the mean** parameter levels among cells within each cell population.

**Table 1.** **A**. Mean of SDs and SD of SDs of AP and Ca^2+^ domain intervals among individual cells in each of the 3 steady state populations that differ with respect to autonomic receptor stimulation; **B**. Mean ± SD of means of AP and Ca^2+^ domain interval in each cell population (A).

| **A. Mean SD ± SD of SDs (ms, individual cells within each population)** |  |  |  |
| --- | --- | --- | --- |
| **AP recordings** | **ISO (n=27)** | **Control (n=78)** | **CCh (n=10)** |
| APFI, SD | 8.43 ± 1.90 | 13.30 ± 5.18 | 99.08 ± 57.28 |
| TTIO, SD | 13.14 ± 4.54 | 23.72 ± 13.33 | 132.78 ± 88.24 |
| APD<sub>90</sub>, SD | 4.52 ± 1.91 | 9.23 ± 4.56 | 78.40 ± 58.07 |
| **Ca<sup>2+</sup> recordings** | **ISO (n=27)** | **control (n=78)** | **CCh (n=10)** |
| CaTFI, SD | 17.52 ± 14.30 | 16.23 ± 11.24 | 86.08 ± 54.28 |
| CaT<sub>90</sub>, SD | 17.31 ± 12.34 | 13.17 ± 7.81 | 30.79 ± 13.35 |
| LCR period, SD | 38.11 ± 24.77 | 40.34 ± 23.90 | 153.31 ± 93.59 |
| **B. Mean ± SD of Means (ms)** |  |  |  |
| **AP recordings** | **ISO (n=27)** | **Control (n=78)** | **CCh (n=10)** |
| APFI, Mean | 244.87 ± 22.13 | 321.57 ± 63.08 | 786.45 ± 374.35 |
| TTIO, Mean | 188.01 ± 18.69 | 239.45 ± 60.45 | 690.44 ± 334.27 |
| APD<sub>90</sub>, Mean | 123.55 ± 14.06 | 169.06 ± 35.37 | 249.72 ± 81.38 |
| **Ca<sup>2+</sup> recordings** | **ISO (n=27)** | **control (n=78)** | **CCh (n=10)** |
| CaTFI, Mean | 357.80 ± 64.94 | 390.64 ± 83.37 | 872.84 ± 284.33 |
| CaT<sub>90</sub>, Mean | 160.76 ± 43.49 | 174.73 ± 42.73 | 289.94 ± 70.24 |
| LCR period, Mean | 302.92 ± 55.65 | 342.68 ± 72.87 | 734.92 ± 191.01 |

Figure 7 A illustrates the distributions of **SDs** of parameters within Ca^2+^ and V_m_ domains **during** APs and, of AP firing intervals in cells listed in Table 1A of each of the three populations of cells that differed in autonomic state: control cells (n=78); cells during super-fusion with ISO (n=27); and in cells super-fused with CCh (n=10). The self-similarity (long-range correlations) of the mean SDs of Ca^2+^ to V_m_ parameter transitions during ignition and recovery phases of APs across the wide range of AP firing intervals induced by the type of autonomic receptor stimulation, or lack thereof, is evident in the self-similarity of their mean **SD** distribution **shapes** (Figure 7A).

**Figure 7.**
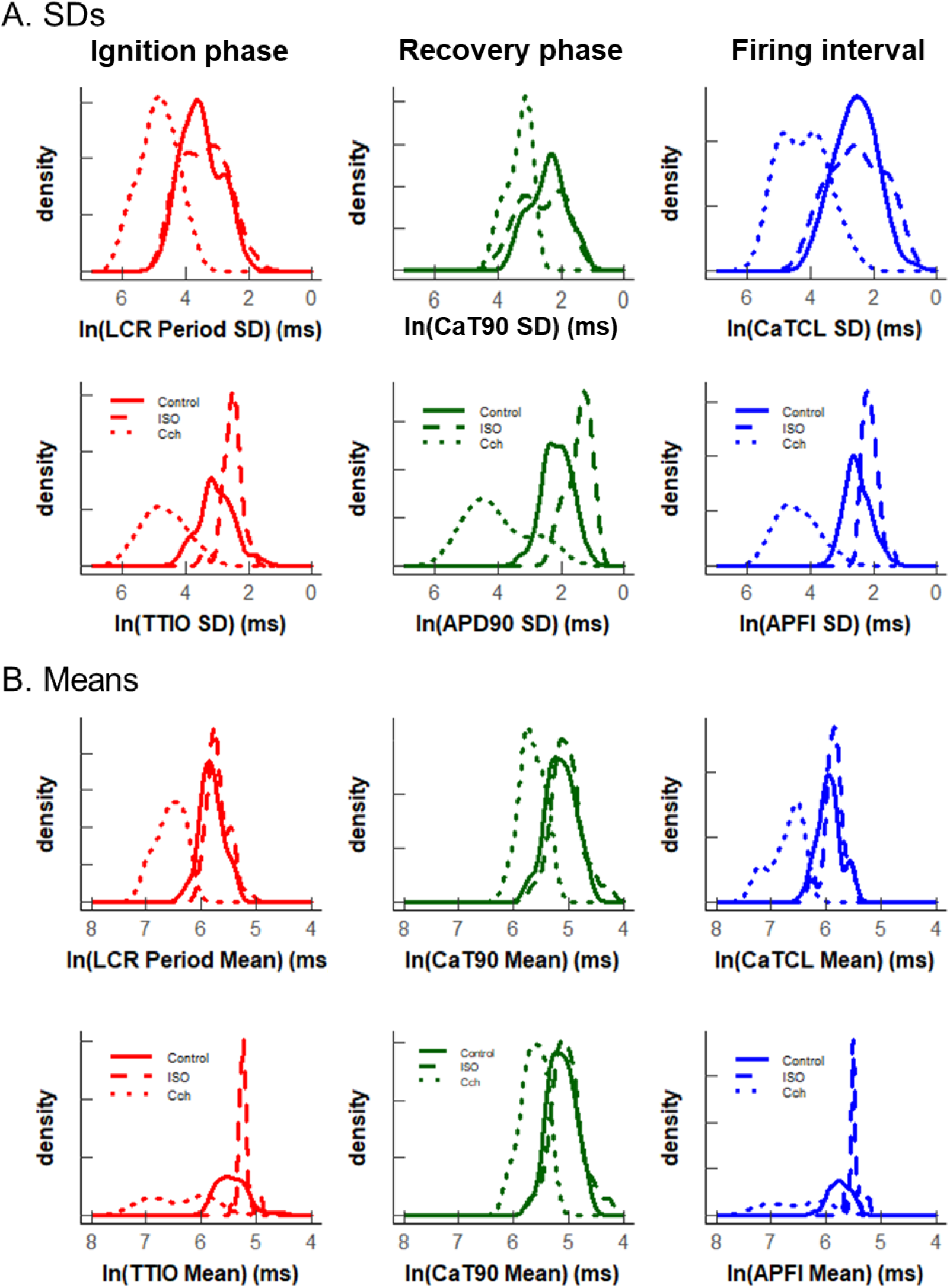
Density distributions of selected parameter means (B) and SDs (A) of M and Ca^2+^ clock functions measured in different cells prior to and during autonomic receptor stimulation across the 3 groups of mean APFI steady states in Table 1. The density distributions are presented as nonparametric kernel estimates of probability density functions (Silverman, 1986), scaled so that the total area within each curve is unity. Both the mean and variability about the mean are concordant with each other across the 3 experimental groups in control and shift concordantly in response to autonomic receptor stimulation in all cells measured.

The distribution of the **means** listed in Table 1B are illustrated in Figure 7B. Note that the **shapes** of the distributions are self-similar to each other across the three different autonomic states. Note also that the shapes of the distribution of the **means** of a given parameter in Figure 7B are similar to the distribution of that parameter’s SDs in Figure 7A (because the interval distribution means stem from the distributions of their SDs). In other terms, the variability in the times at which parameters occur within a time series (their parameter SDs) determines what the mean interval of events in the time series will be.

Also note in Figure 7, that compared to cells not super-fused with an autonomic receptor agonist (control cells) and those super-fused with ISO, the shapes of the distributions of cells super-fused with CCh are broad, indicating marked variability among CCh cells within the parameter distributions of both interval means and interval SDs.

Figure 8A illustrates the self-similarity of V_m_ parameter means and SDs to Ca^2+^ means and SDs across autonomic states. Heat maps of the long-range correlations among V_m_ and Ca^2+^ parameter means are shown in Supplementary Figure S3. The long-range correlations (self-similarity) between the means of the means and means of SDs given in table 1 is shown in Figure 8B.

**Figure 8.**
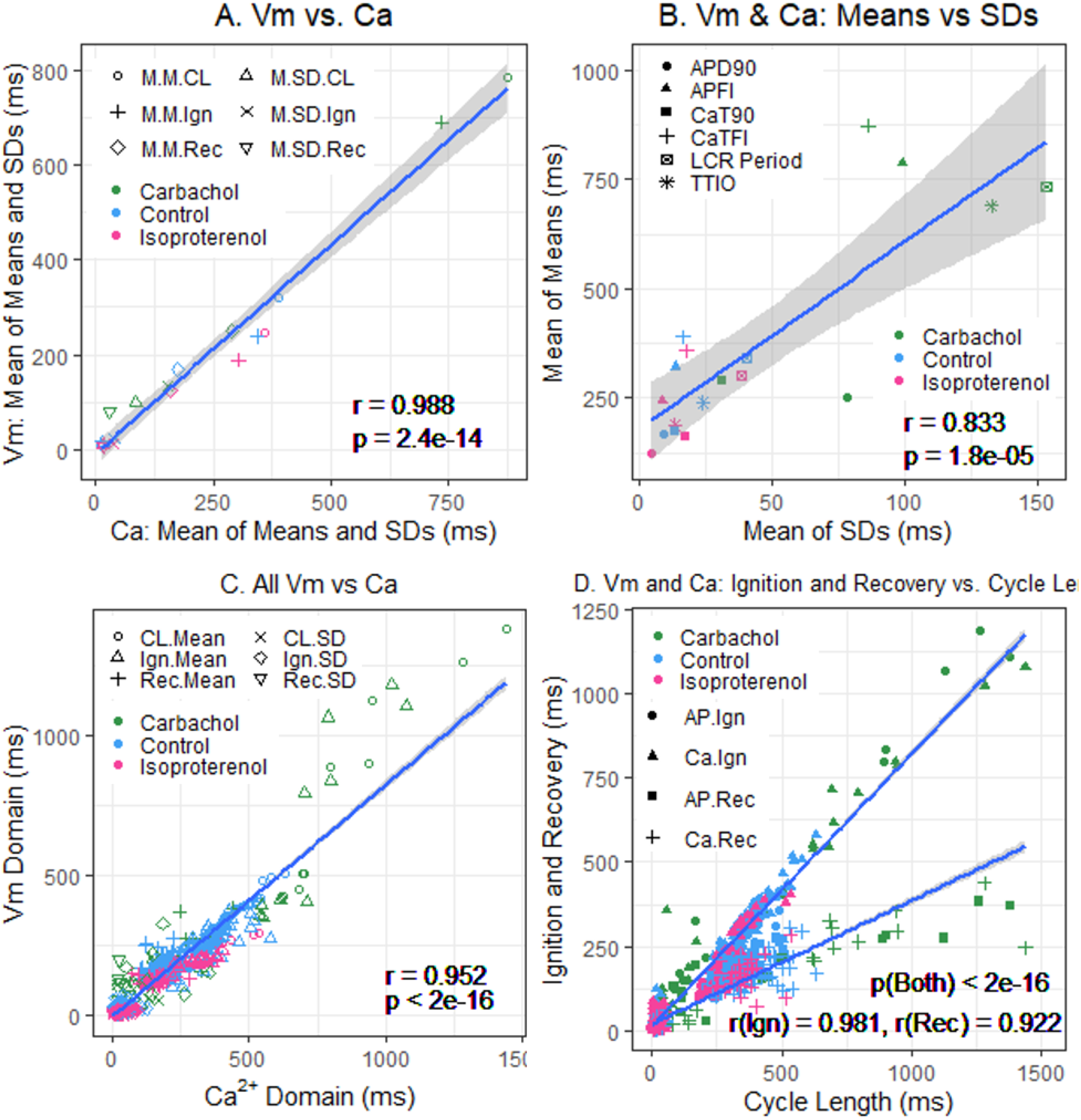
A. Plot of mean of Means and SDs of Vm vs. Ca^2+^ from Table 1; B. Mean of means in Table 1 vs. mean of the SDs in Table 1; C. Means and SDs of Vm parameters vs. Means and SDs of corresponding Ca^2+^ parameters for all 230 cells; D. Means and SDs of Ignition and Recovery vs. Means and SDs of Cycle Length for all 230 cells (Note: In Panel A, M.M = mean of means; M.SD = mean of SDs. CL= cycle length, Ign = Ignition, and Rec = Recovery).

### Self-similarity of Ca^2+^ and V_m_ parameters among <u>all individual cells</u> within and among the three autonomic states

Two by two correlations of V_m_ and Ca^2+^ parameter **means** and **SDs** of all (230) cells that comprised the three different cell populations in Figure 7 and Table 1 are highly significant (Table 2).

**Table 2.**
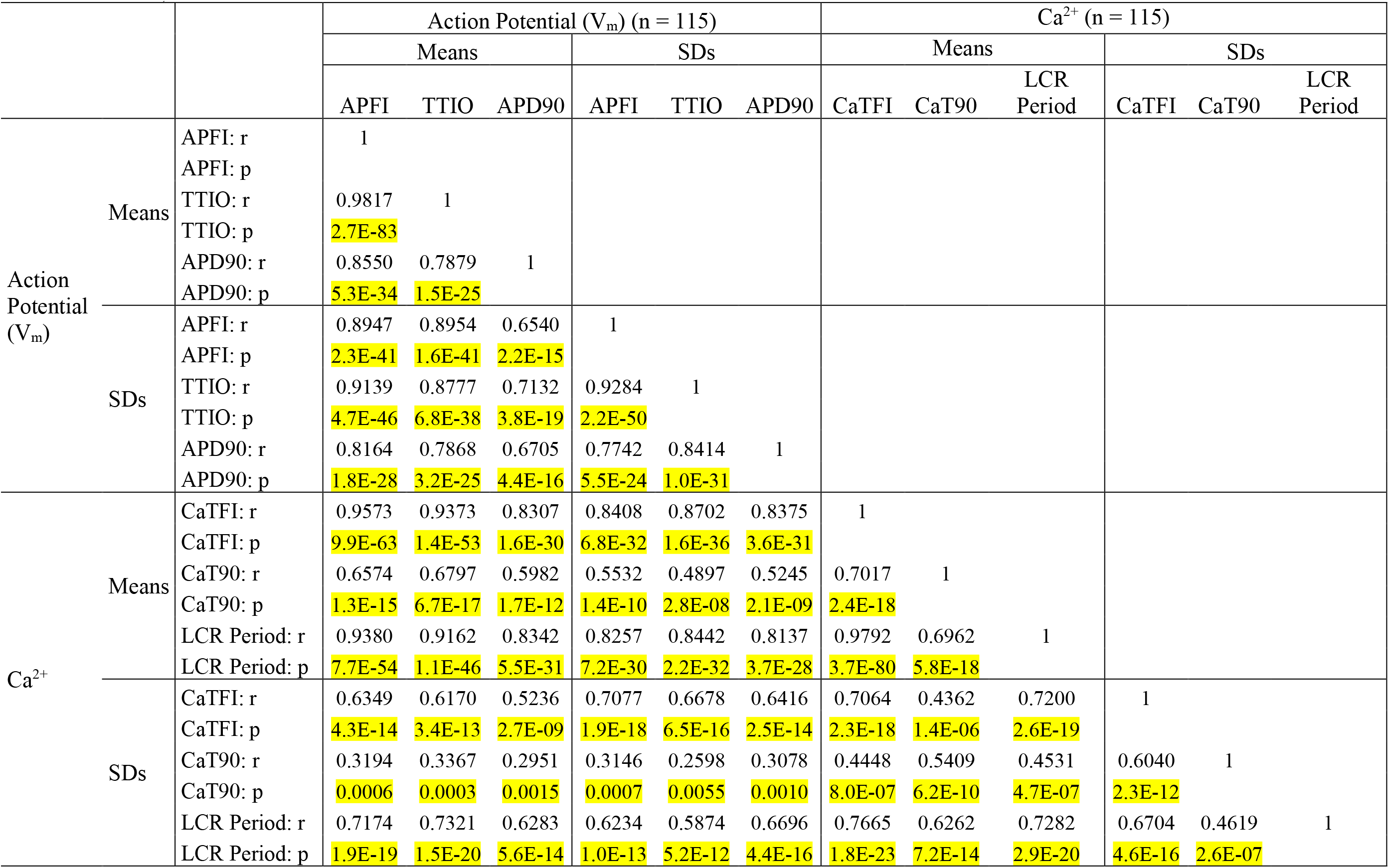
Correlation matrices of Vm or Ca^2+^ parameters measured in different cells within 3 different autonomic states (summary data listed in Table 1).

Figure 9 shows ln-ln plots of the distributions of means and SDs of Ca^2+^ and V_m_ parameters of all cells across the three autonomic states. The piecewise linear fit of the data in each panel is largely driven by the data from cells super-fused with CCh that manifested broad interval distributions and high mean APFIs in Figures 4 to 7. Figure 8C shows that the means and SDs of V_m_ parameters measured during APs in individual cells (n=115) are self-similar to Ca^2+^ means and SDs measured during AP in other individual cells (n=115) across the three autonomic states.

**Figure 9.**
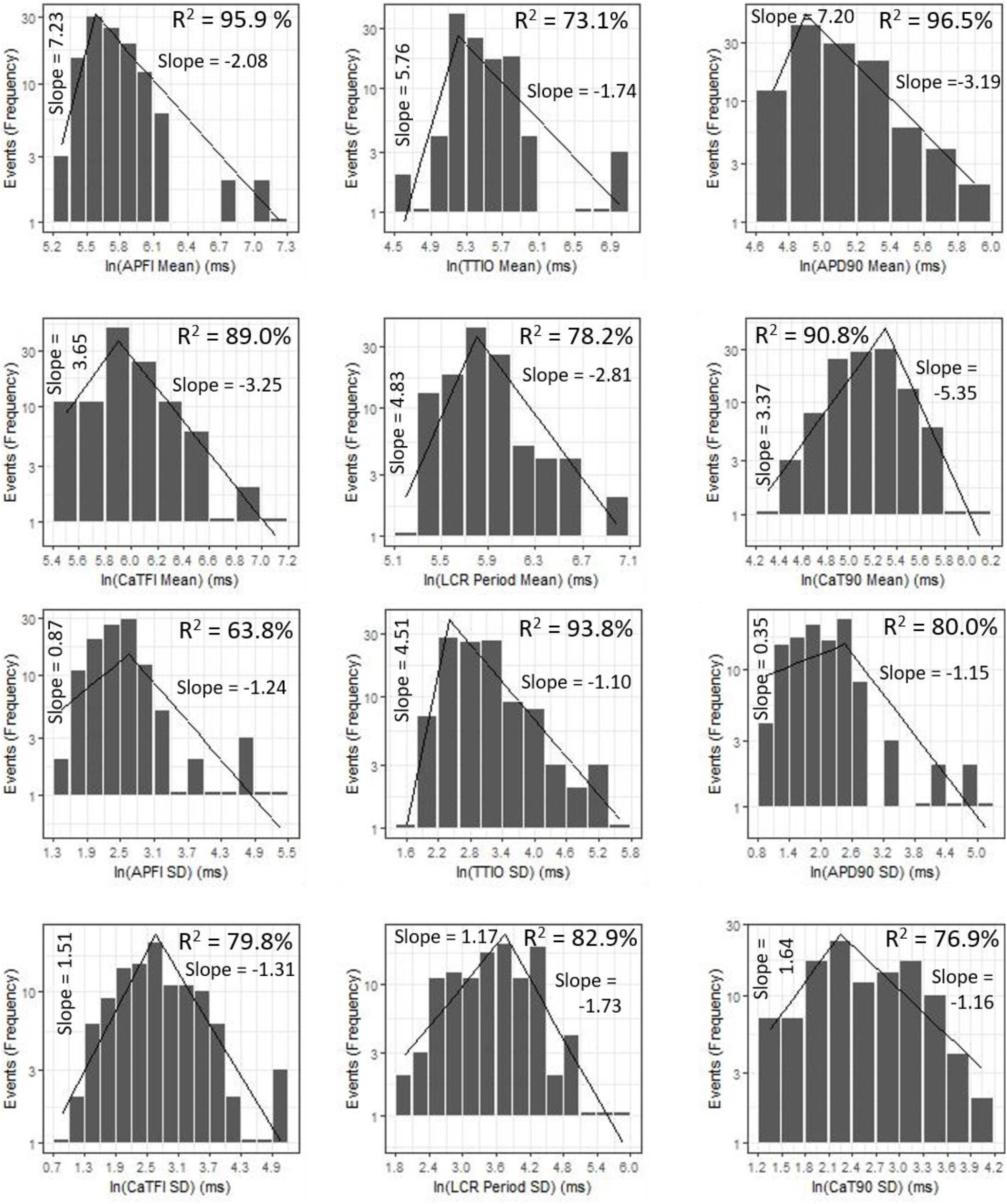
Distributions of SDs and Means over all groups illustrating self-similarity (Fractal-like behavior). The regression lines are fit as a single piecewise linear model with a join point at the center of the interval with the highest frequency.

Although, as noted above, the V_m_ and Ca^2+^ parameters within and among cells during CCh super-fusion were more broadly distributed than those during control or during ISO, the correlations between V_m_ and Ca^2+^ parameters among all cells **within** the CCh super-fused population of cells were extremely strong for most parameters (Supplementary Figure S3). Weaker but still significant correlations between times to 90% recovery and other variables are observed in CCh super-fused cells (likely because times to 90% are the most difficult parameters in the data set to measure accurately).

Figure 8D shows that V_m_ and Ca^2+^ parameters measured during APs are self-similar to AP firing interval means and SDs across the three autonomic states.

### Correlation of Ca^2+^ and V_m_ domain parameter means in individual cells to their SDs within and across autonomic states

The relationship between mean AP firing rate and its SD is known to be non-linear (Monfredi et al., 2014). Although the relationships of **all** Ca^2+^ and V_m_ parameter **means** relative to their corresponding **SDs** measured in the combined set of data derived from different populations of cells in control or during ISO or CCh super-fusion are non-linear (Figure 10, A-C), the ln-ln plots of this combined data (Figure 10, Panels D-F), however, are linear, indicating their self-similarity across **all** 230 cells that differed by autonomic state.

**Figure 10:**
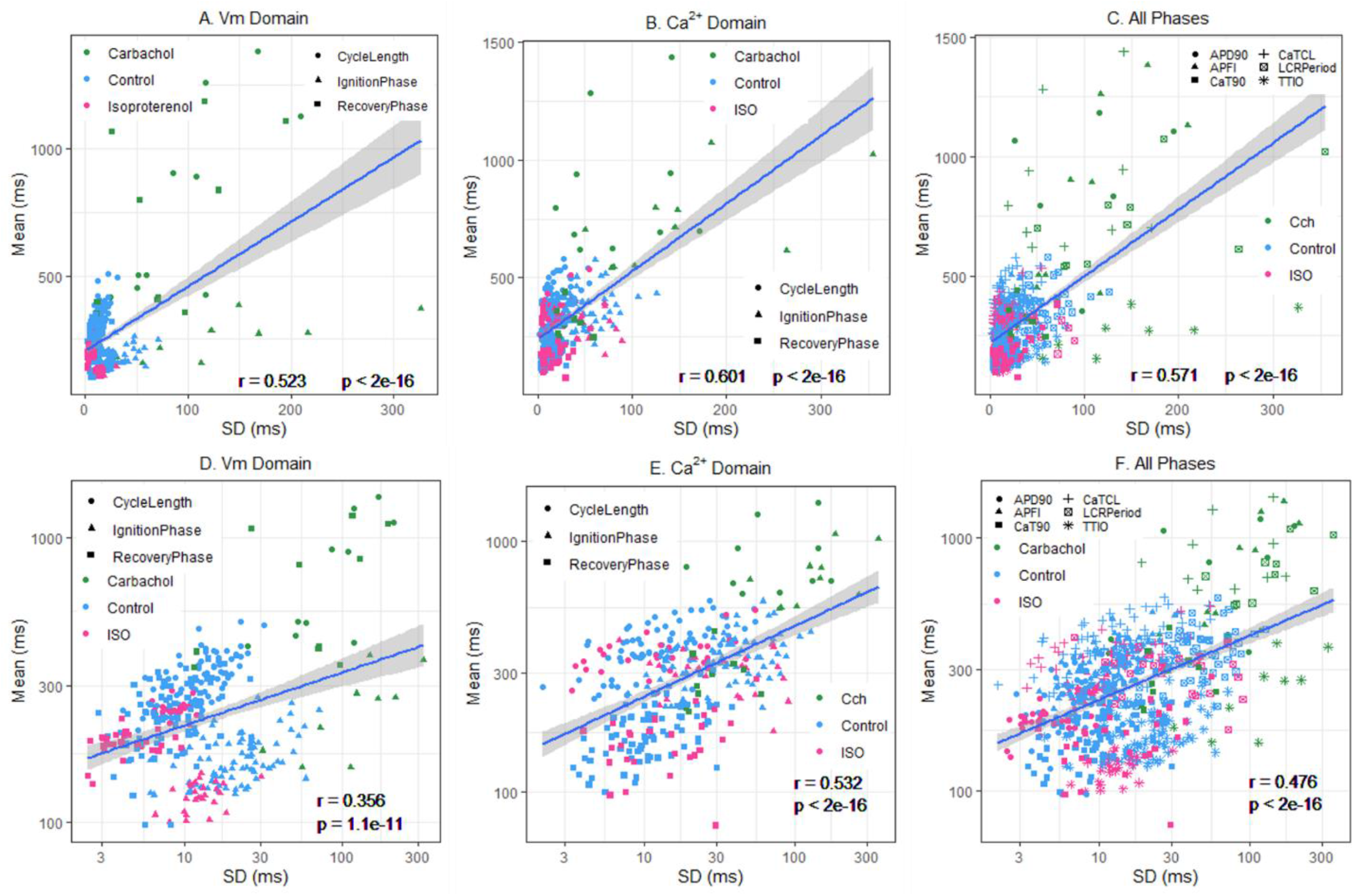
Means vs SDs for all cells on a linear scale for A. V_m_, B. Ca^2+^, C. V_m_ and Ca^2+^; and using a logarithmic scale for D. V_m_, E. Ca, F. V_m_ and Ca^2+^.

### Principal component analyses

Next, we employed principal component analyses to determine whether the self-similarity of parametric measures within the entire data set of variables could be summarized by a smaller set of principal components that contain most of the information in all the variables. PCs are linear combinations of the original variables, and each PC is statistically independent of the others: the first PC explains as much of the total variability in the data as possible, the second PC as much of the remaining variability, and so on. Highly self-similar parameters within the complete data set are explained by the sum of the first few PCs.

In a PC analysis of SDs of the six variables (3 in the V_m_ domain and 3 in the Ca^2+^ domain) the first 3 PCs explained 91.4% of the total variation within the entire SD data set (Supplementary Table S2, Figure 11A). Similarly, in a PC analysis of the means of all six measured variables means (3 in the V_m_ domain and 2 in the Ca^2+^ domain) in a PC analysis, the first 2 PCs explained 92.8% of the variability in the data set of all 6 means (Supplementary Table S2, Figure 11B). Finally, in a PC analysis of all 12 variables (6 means and 6 SDs), the first 3 PCs explained 88% of the variability within the total (means plus SDs) data set (Supplementary Table S2, Figure 11C).

**Figure 11:**
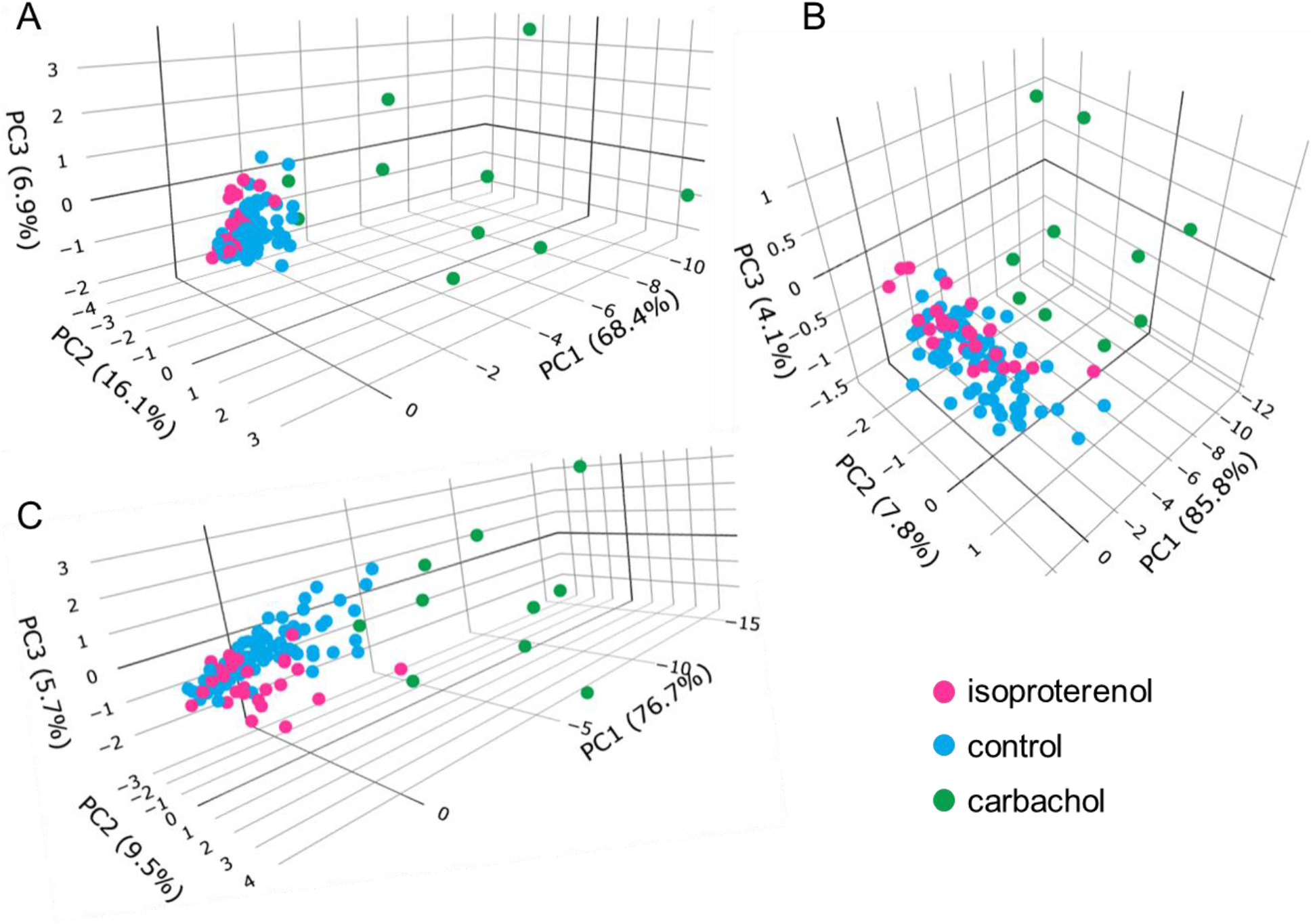
Plots of Principal Components: A) SDs only; B) Means only; C) Means and SDs.

Because a smaller set of PCs can explain a substantial proportion of the total variability in each set of V_m_ **or** Ca^2+^ domain means, SDs, and means and SDs, means that these distributions of V_m_ and Ca^2+^ parameters measured during an AP, and AP firing intervals in control cells and different cells super-fused with ISO or CCh are each self-similar to each other. In other terms, Ca^2+^ and V_m_ domain functions operative within the SAN cells coupled-clock system manifest self-similar-scale free characteristics i.e., are kinetic fractals of each other, across the entire physiologic range of AP firing intervals.

### Numerical model simulations of APFIV, major ion currents, and Ca^2+^

Variability of V_m_ and Ca^2+^ parameters measured experimentally in cells within and across autonomic states is linked to the respective variabilities of clock molecular availability to respond to V_m_ and Ca^2+^ cues (Figure 2) that cannot be directly measured experimentally during AP firing. To gain further insight into the variability of these biophysical mechanisms, we performed numerical modeling simulations. The APFI variability was generated by SAN cell model described as a stochastic dynamical system, i.e., a dynamical system (deterministic Maltsev-Lakatta model (V. A. Maltsev & Lakatta, 2009)) subjected to the effects of noise current, I_per_ (see Methods and Supplementary Material for details). I_per_ amplitude was tuned for the model APFIV to match that measured experimentally under respective experimental conditions. We investigated two scenarios of noise generation: when I_per_ was added to I_tot_ or when I_per_ was added to Ca^2+^ release flux current in each of the three autonomic states: (i) basal AP firing, (ii) ISO 100 nM, and (iii) CCh (100 nM). Variability of 6 major currents was simulated and analyzed: I_f_, I_NCX_, I_Kr_, I_CaL_, I_CaT_ and I_KACh_. Variability of [Ca] under cell membrane was also simulated during the three autonomic states. For all items we measured variability of their peak amplitudes and amplitudes at −40 mV during DD.

Model simulation results are presented in Figure 12, with their numerical values given in supplementary Tables S6 and S7. Regardless of the type of noise generation (via Ca^2+^ or I_tot_), it affected the variability of ion currents and Ca^2+^ the same way and the predicted variabilities for many parameters differed substantially from that of APFI:

1. I_f_ variability was substantial: in the basal state and in ISO I_f_ variability was similar to or larger than APFI variability; The variability of I_f_ decreased in CCh.
2. I_NCX_ variability was also substantial: at −40 mV it was substantially larger than that of APFI variability (except in CCh when I_per_ was added to I_tot_); variability of I_NCX_ peak amplitude (negative, during AP upstroke) was similar to that of APFI in the basal state and ISO, but became reduced in CCh.
3. I_Kr_ variability was substantially less than APFI variability under all conditions.
4. I_CaT_ variability was the largest among ion currents, being similar to that of I_NCX_, in CCh when I_per_ was added to Ca^2+^ release.
5. Peak I_CaL_ variability was always less than that of APFI variability; at −40 mV it was greater in ISO than in basal state and CCh.
6. Variability of Ca^2+^ release flux in the basal state and ISO at −40 mV was greater than or similar to APFI variability; in CCh variability of Ca^2+^ release flux was less than that of APFI.

**Figure 12.**
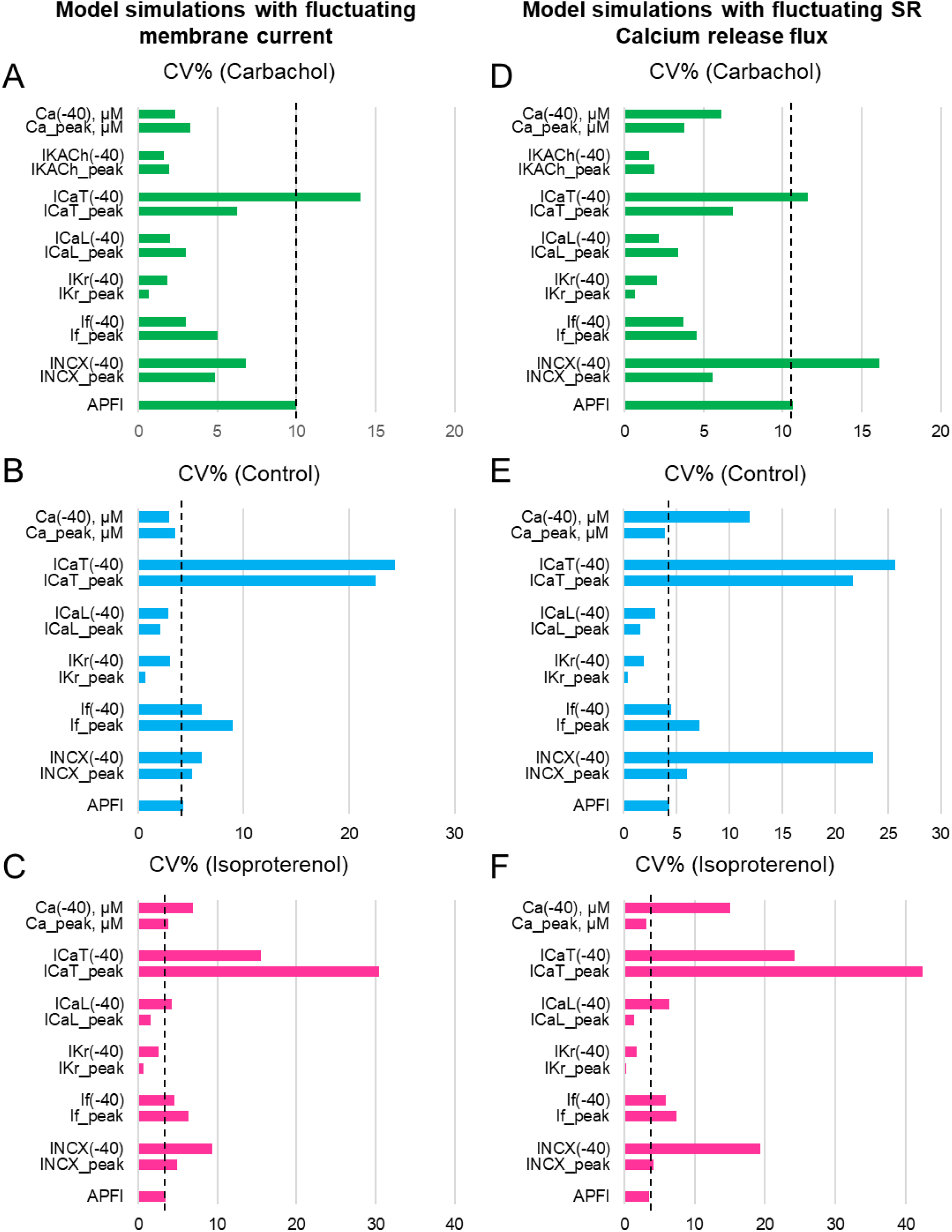
Results of analysis of coefficient of variation (CV) of major ion currents I_f_, I_Kr_, I_CaL_, I_CaT_, I_KACh_, and Ca^2+^ simulated by Maltsev-Lakatta coupled-clock model with noise current (I_per_) added to either total current I_tot_ (panels A~C) or Ca release flux (panels D~F).

Some components exhibited power law behavior over a wide range of APFI over all conditions tested (Figure 13).

**Figure 13.**
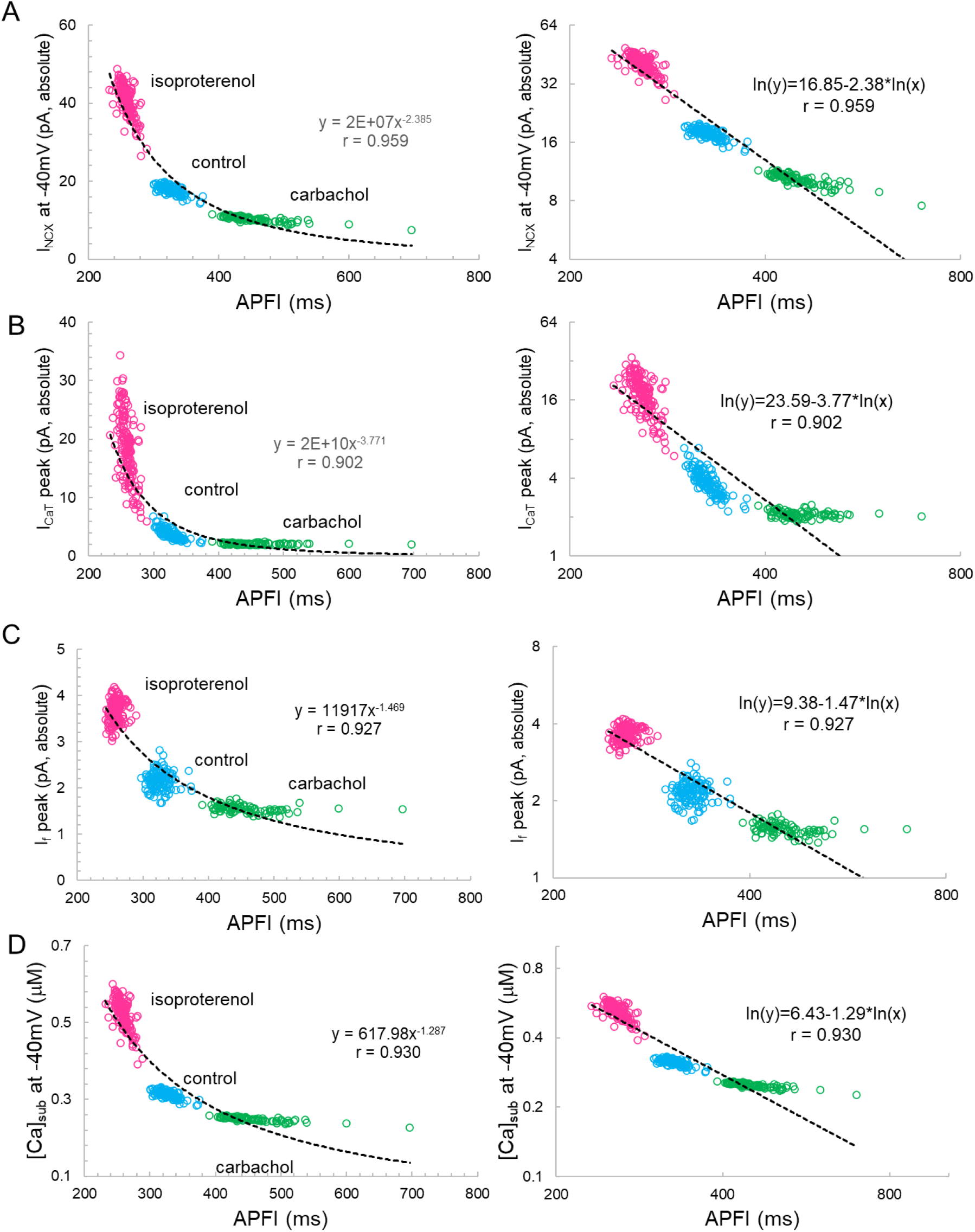
I_NCX_(A), I_CaT_(B), I_f_,C), and [Ca]_sub_ (D) exhibited power law behavior (a linear dependence in the ln-ln plot here) over a wide range of APFI over all conditions tested. Noise (I_per_) was added to I_tot_ in Maltsev-Lakatta coupled-clock model. Similar dependencies were found when I_per_ was added to Ca release flux (not shown).

We next determined whether self-similarity across autonomic states observed for experimental data during the ignition phase is also applied to simulated ion currents or Ca^2+^ data during this time of the cycle, i.e. at −40 mV. To this end, we applied the statistical tests utilized for experimental data to simulated data (for the scenario when I_per_ was added to I_tot_).

Simulated ion currents and Ca^2+^ amplitudes during AP ignition (−40 mV) across the three autonomic states are self-similar to each other, strongly correlated to each other, as were experimentally measured parameters (Table 2). These two-by-two correlations of all the simulated components are listed in Supplementary Table S3. Selected examples of these correlations are shown in Figure 14. As I_NCX_ is fully determined by V_m_ and Ca^2+^, at a fixed voltage (−40 mV) it is fully determined by only Ca^2+^. That is why we have 100% correlation of I_NCX_ and Ca^2+^. I_CaT_ strongly correlated with Ca^2+^ variations, because the stronger Ca^2+^ signal is linked to the higher DD rate and hence stronger (time-dependent) activation of I_CaT_. Surprisingly I_KACh_ amplitude was also highly correlated with Ca variations.

**Figure 14.**
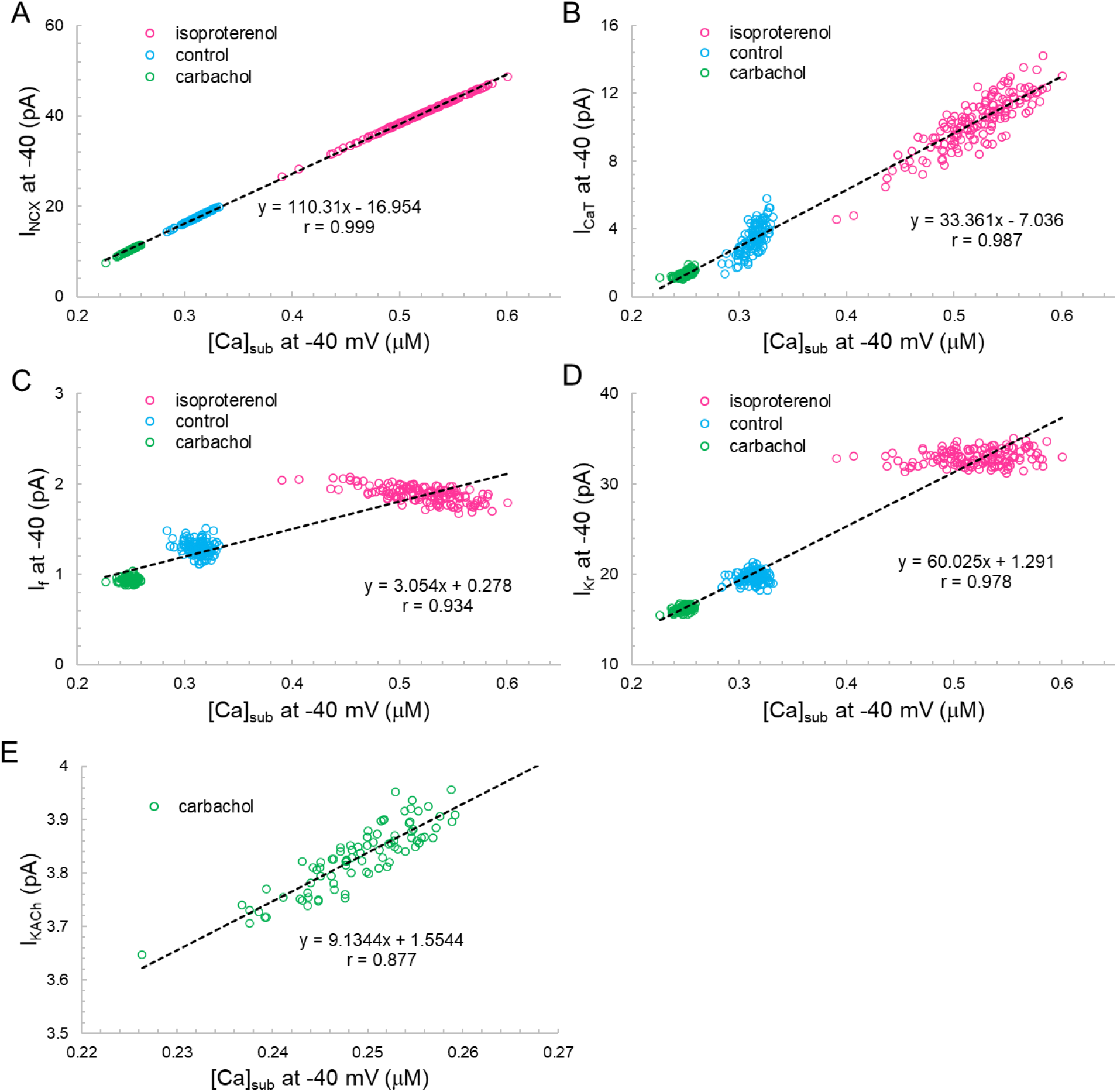
The correlation between I_NCX_(A), I_CaT_(B), I_f_, (C) I_Kr_(D) or I_KACh_(E) vs. [Ca]_Sub_ at −40mV. Note that I_KACh_ only presents during CCh.

Variations in I_f_ and I_Kr_ at −40 mV did not depend on variations of Ca^2+^, but their mean values strongly depended on Ca^2+^ across the autonomic states. I_f_ activation and I_Kr_ deactivation are early DD mechanisms and do not seem to interplay with Ca^2+^ at the ignition onset at −40 mV in a given cycle.

Figure 15A illustrates the Poincare plots of many simulated parameters (APFI and TTIO; I_NCX_, I_CaL_, I_Kr_, [Ca], I_CaL_, I_KACh_, and I_f_, all at −40 mV). Although the range of absolute values of these simulated parameters substantially vary, all simulated parameters are self-similar, i.e. are fit by a single line (r=0.998) with a slope of unity, resembling Poincare relationship of experimentally measured AP parameters (Figures 1 and 4).

**Figure 15.**
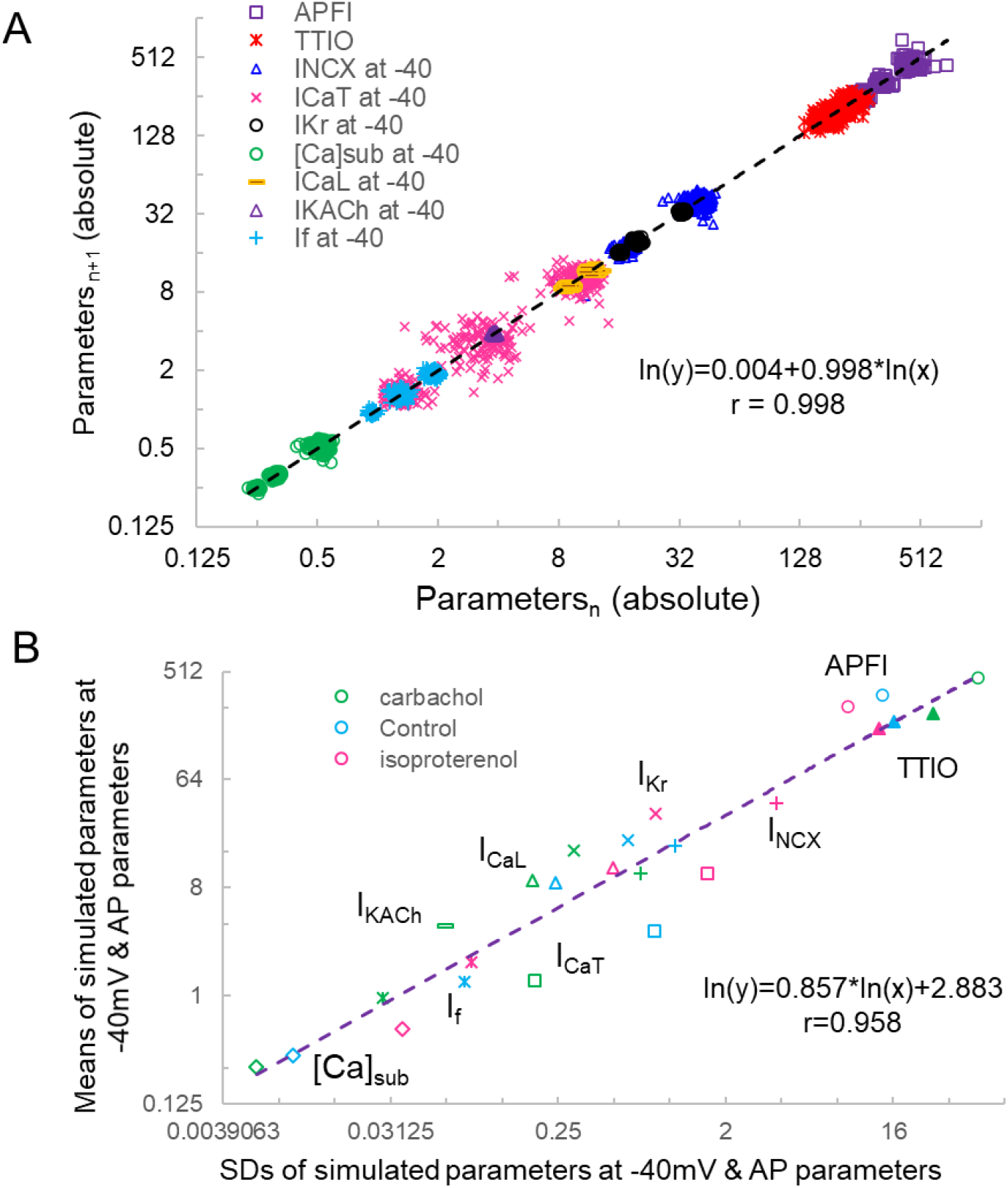
Poincaré plots of the simulated I_NCX_, I_CaT_, I_CaL_, I_Kr_, I_KACh_, I_f_, [Ca]_Sub_ at −40mV, APFI and TTIO from 1 model cell during ISO, CCh or at basal condition in ln-ln plot (A) and the correlation between the means and SDs of these parameters (B). Note that I_KACh_ only presents during Carbochol.

Figure 15B illustrates ln-ln plots of the relationships of the means of simulated components to their SDs. Note that this relationship follows power law behavior just as did the relationship of experimentally measured means for AP parameters vs their SDs (Figure 10).

Finally, in PC analyses of the simulated parameters in Figure 15B the first 2 PCs accounted for 94% of the variation in the eight variables (Supplementary Figure S4).

## Discussion

We measured membrane potential and Ca^2+^ times to onsets of AP ignition during diastole, and times to 90% recovery during APs, and AP firing intervals in 3 populations of single, isolated rabbit SAN cells that differed with respect to autonomic input: those in which CRs were stimulated by CCh; those in which βARs were stimulated with ISO; and in untreated (control cells). **Absolute values** of times to AP ignition onsets and to 90% recovery intervals in Ca^2+^ and V_m_ domains during APs and AP firing intervals differed within and among individual cells in cell populations and different autonomic states, and differed **markedly** among cells of the three populations of cells with differential autonomic receptor stimulation. Our novel finding is that although differing markedly in absolute values, Ca^2+^ and V_m_ parameters were self-similar to each other during APs and self-similar to AP firing intervals, not only within and among different cells **within** each of the three populations of cells studied, but, remarkably, **among all** cells, regardless of the autonomic receptor stimulation profile. Thus, Ca^2+^ and V_m_ domain kinetic transitions (intervals) during APs, individual AP firing interval and mean AP firing intervals within AP time series manifest long-range correlations (self-similar scale-free correlations, i.e., obey power law) across the entire broad range of AP firing intervals, regardless of whether autonomic receptors of these cells are stimulated or not, and regardless of the type of autonomic receptor stimulation.

The degree to which molecular activation states within each clock and between clocks are synchronized during APs determines when the next AP will occur, i.e. the AP firing interval variability and mean AP firing intervals within a cell and across the entire population of single SAN cells studied: the higher the degree of order (self-organized activation of clock molecules) the more ordered and less variable the aggregate of kinetic functions, the least variability of AP firing intervals and the shorter the **mean** AP firing interval; vice-versa, the lower the degree of order among clock molecular activation states the lower the aggregate synchronization among clock molecular functions, the greater the variability of AP firing intervals, the longer the **mean** AP cycle interval.

Self-similar or fractal-like beating rate variability among cardiac cells in culture has been previously identified in a number of studies **but only when cells were confluent**, or electrically connected to each other. This behavior has been attributed to influences of tonic or phasic resetting of membrane potential, or to mechanical factors via cell-to-cell connections (Clay & DeHaan, 1979; Jongsma, Tsjernina, & de Bruijne, 1983; Kucera et al., 2000). Our novel observation is selfsimilarity of V_m_ and Ca^2+^ domain intervals during APs and AP firing intervals across diverse populations of single SAN ells that were not physically connected to each other.

Thus, self-similar distributions of order that have been demonstrated to occur in other instances throughout nature (Bak, 1999), also exist within SAN cell coupled-clock system functions. We interpret this power law behavior of SAN cell functions to result from concordant gradations of self-organized order (synchronization) of clock molecular activation across the entire physiologic range of AP firing intervals.

### Clock molecular activation cues

Voltage, time, Ca^2+^, cAMP signaling, and PKA and CaMKII-dependent clock protein phosphorylation, are the cues regulate the activation kinetics of molecules that control pacemaker functions in single SAN cells (Supplementary Figure S2) (Lakatta et al., 2003; Lakatta et al., 2010; Lakatta et al., 2006; Lakatta et al., 2008; V. A. Maltsev & Lakatta, 2008; Yaniv et al., 2015). Some coupled-clock system proteins are activated by Ca^2 +^ e.g. SERCA 2; others by V_m_ and cAMP binding, e.g. HCN channels (DiFrancesco & Tortora, 1991) and other cyclic nucleotide-regulated channels; or by Ca^2+^ **and** V_m_, e.g. NCX, or by phosphorylation **and** Ca^2+^ e.g. phospholamban and Ryanodine Receptors and AC type 8; while the activation states of still other coupled-clock system proteins are modulated by V_m_, Ca^2+^ **and** phosphorylation, e.g. L type and some K^+^ channels.

Both voltage and Ca^2+^ activation cues oscillate in amplitude throughout each AP cycle and command rapid responses from clock molecules. The degree to which activation status of molecules of a given species is synchronized at any given time following the prior AP determines the ensemble response of that molecular species to its activation cues. It is well documented that following a synchronizing event, e.g. the occurrence of an AP, activation states of molecules underlie AP cycle transition through variably inactivated states, altering the availability to respond to a subsequent activation cue. Our new concept of synchronization of functional cues is based on the idea that the coupled-clock system inheres (inevitably) some degree of disorder that stems from its key constituent proteins operating (stochastically switching) intrinsically within their conformational flexibilities and heterogeneity. The balance of order/disorder is linked to molecule interactions (i.e. effectiveness of their respective cues) that allows them to operate cooperatively as an ensemble or system with various degree of synchronization (i.e. order) that is reflected in respective variability of the output function of the system, i.e. APFI variability in our case.

Thus, we interpret the experimentally measured concordant behavior of surface membrane and Ca^2+^ regulatory functions during AP cycles across the entire physiologic range of AP cycles to reflect a concordance in the degrees of activation of molecules that drive these regulatory functions. Importantly Ca^2+^ and V_m_ cues **not only** regulate the synchronization clock of molecular activation states but are also **regulated by** the degree of synchronized activation of molecules determined by these cues (recursion). Because membrane and Ca^2+^ clocks become coupled in the context of the electrochemical signal that waxes and wines to cause the AP cycle, the extent of self-organized molecular activation **within** each clock indirectly affects self-organization of molecular activation of the **other** clock operating within the coupled-clock system. And because scaling of **mean** APFIs among all cells is self-similar to APFI **variability** among cells, AP firing variability and mean APFI are determinants of the Ca^2+^ and V_m_ cues that determine kinetic intervals during an APs: a recursive, feed-forward process.

### Mean APFI and APFIV are not only <u>regulated by</u>, but also <u>regulate</u> the degree to which clock molecular functions are synchronized

Changes in Ca^2+^ and V_m_ cues during an AP, not only determine the characteristics of that AP, but also determine when the next AP will occur, and the mean AP firing interval within an AP time-series.

A prolongation of the mean APFIs, itself, contributes to the concurrent increase in the AP firing interval variability at long mean APFI: because an increase in mean APFI reduces net Ca^2+^ influx, and indirectly reduces Ca^2+/^ cAMKII-AC-dependent phosphorylation of Ca^2+^ cycling proteins, reducing the SR Ca^2+^ cycling kinetics and increasing the variability of LCR periods.

Characteristics of the AP that are determined by availability of M clock molecules to respond to a change in membrane potential, both directly and indirectly entrain the Ca^2+^ and M clock activation: As the mean AP interval shortens, less time elapses between APs, and therefore at shorter intervals less time is required than at longer intervals for molecules to retain (remember) the synchronizing influences imparted by the preceding AP. This causes the relationship of mean APFI to APFIV of isolated SAN cells to be non-linear (Figure 5 and Supplementary Figure S2), as originally demonstrated by ZaZa (Zaza & Lombardi, 2001) and later by Monfredi et al (Monfredi et al., 2014). Conversely, as time following a prior AP increases, the effectiveness of the Ca^2+^ activation cue, itself, wanes, because the cell Ca^2+^ level and SR Ca^2+^ load become reduced, due to time-dependent Ca^2+^ efflux from the cell. We may speculate, therefore, that during long AP cycles, fewer molecules of some molecular species are available to respond to Ca^2+^ activation cues.

Gradations of self-organized molecular activation within and between clocks, regulate the APFI rhythm i.e., the APFI variability. In other terms, the average APFI, kinetics of the AP, AP-triggered Ca^2+^-transient, LCR periods and diastolic depolarization kinetics, and beat-to-beat variability of these parameters measured in the present study are readouts of the relative extents to which of clock molecules become activated and the degree to which the clocks are coupled. When the degree to which Ca^2+^ and M clocks kinetics are coupled or synchronized is low, the AP firing rate is slow, and AP firing interval variability is high, e.g., during CRs. Conversely when the degree of coupling or synchronization of the Ca^2+^ and V_m_ kinetics of the two clocks is high e.g., during βAR stimulation, AP firing is rapid and AP firing interval variability is low.

### So, what factors affect the degree of synchronization of clock molecules?

Concordant degrees of self-similar synchronization of M and Ca^2+^ clock kinetic functions reflect concordant gradations of activation states of specific molecules that govern these functions and how these cues change throughout an AP cycle.

### AP firing rate and rhythm synchronization of clock molecules

The AP that emerges from the diastolic ignition events is, itself, the most potent integrator or synchronizer, not only of surface membrane electrogenic molecules, but also of Ca^2+^-clock functions: a synchronized global cytosolic Ca^2+^ transient that ensues following synchronous activation of voltage-dependent L-type Ca^2+^ channels is created by synchronized Ca^2+^-induced, Ca^2+^ release from SR via ryanodine receptor activation. (Lakatta, 2004; Song, Sham, Stern, Lakatta, & Cheng, 1998; Wang, Song, Lakatta, & Cheng, 2001; Zhou et al., 2009).

The efficacy of V_m_ and Ca^2+^ activation cues that oscillate as electrochemical signal that underlies the V_m_ change during AP cycle varies with the AP cycle interval or period: shorter periods (i.e., faster AP firing rates or shorter APFIs) are more effective than longer periods (i.e., slower AP firing rates or longer APFIs), because during very long AP cycles, Ca^2+^ activation states of some molecules become more unsynchronized. At very short times following a large voltage oscillation (i.e., an AP), many molecules of a given molecular species in relatively inactivated state may not optimally respond to activation cues (e.g., impaired excitability/non-excitability). As the time following a prior activation increases, although a sub-population of molecules of given species may regain full ability to respond to activation cues, substantial variability in the activation status of other molecules of that species still may exist, limiting the number of molecules that can respond to (be recruited by) an activation cue. Our results provide novel clues to the cellular basis for the observation that an AP occurrence, itself, influences the range of APFIs that immediately follow it (Nolasco & Dahlen, 1968). The AP, itself, indirectly affects all Ca^2+^-clock functions because it regulates net cell Ca^2+^ balance. Functions of M-clock molecules that underlie the generation of an AP *indirectly* regulate the availability for SR Ca^2+^ cycling by modulation of the level of cell Ca^2+^, the SR “oscillatory substrate”. Thus, M clock functions also *indirectly* regulate LCR periods and sizes via their impact on the “steady state” intracellular Ca^2+^ level. When the average interval between APs becomes prolonged, a reduction net Ca^2+^ influx into: efflux from the cell (Lakatta, 2004) reduces the cytosolic [Ca^2+^], the rate of Ca^2+^ pumping into SR, and the SR Ca^2+^ load. These reductions, in turn, prolong the average time from the prior AP occurrence for spontaneous local diastolic ryanodine receptor activation to occur within SAN cell local micro-domains; the randomness of spontaneous local diastolic ryanodine receptor activation occurring within these micro-domains also increases, broadening the distribution of LCR periods and shifting these to longer times at long AP cycle (Figure 4).

Thus, the degree of variability in activation states of M and Ca^2+^ clock molecules that emerges over time following their synchronization by the prior AP is implicated in the cycle length dependence of variability of Ca^2+^ and M clock functions measured here (Table 1, Figures 3–5). Heartbeat variability in *vivo*, and AP firing interval variability (APFIV) of isolated SAN cells in *vitro* indicate that, neither autonomic input to SAN cells, nor functions intrinsic to the SAN cell coupled-clock system, respectively, achieve a steady-state from one beat to the next.

### Ca^2+^-dependent synchronization of clock molecules

The local [Ca^2+^], itself, also serves as a powerful synchronizer of clock molecular function: ordered/disordered Ca^2+^ regulation has been recently reported for ryanodine receptor -mediated Ca^2+^ releases (A. V. Maltsev, Stern, & Maltsev, 2019).

Studies in **permeabilized** SAN cells, in which Ca^2+^-clock function is preserved, but M clock function is abolished, and therefore AP’s cannot occur and do not influence LCR periodicity, clearly demonstrate that: in a fixed, physiologic, free [Ca^2+^], LCR occurrences are random when the free [Ca^2+^] is low; and that LCR periodicity emerges as the free [Ca^2+^] in the system is increased, due in part, to an increase in the Ca^2+^ charge of the SR capacitor (S. Sirenko et al., 2013). The intracellular concentration of the oscillatory substrate, Ca^2+^, itself is regulated, in part, by the SAN cell transmembrane Na^+^ gradient and membrane potential (S. Sirenko et al., 2016; S. Sirenko et al., 2013).

### cAMP activation or phosphorylation of clock proteins modulate the synchronization of and response to activation cues

Autonomic receptor stimulation modulates both the activation cues and responses of clock molecules to these cues. The impact of autonomic receptor signaling on the effectiveness of clock-coupling occurs over several AP cycles, and is reflected in time-dependent transitions in the AP firing rate and rhythm. The kinetics and stoichiometry of increases in PKA activity in response to gradations in βAR stimulation predict the kinetics and stoichiometry of concurrent time-dependent increases in AP firing rate (Yaniv, Lyashkov, et al., 2014). Prior studies (Lyashkov et al., 2009; D. Yang et al., 2012) have demonstrated that gradations in the phosphorylation status of phospholamban at Ser^16^ across the 3 autonomic state mean APFI’s of cell populations in the present study (Supplementary Figure S3) strikingly resemble gradations of the means of APFIs and APFIVs observed across these autonomic states measured in the present study (Table 1).

The extent to which clock molecules respond to Ca^2+^ or V_m_ activation cues during an AP is **modulated** by βARs or CRs dependent-phosphorylation of many of the **same** proteins that drive SAN cell automaticity in the absence of autonomic receptor activation. This βARs or CRs impacts on the memory of the extent to which clock molecules activation had been synchronized during the prior AP. βARs or CRs modulation has two facets: (1) a direct effect, due to cAMP or phosphorylation-dependent activation of clock proteins and (2); an indirect effect by altering the APFI, which alters the cell Ca^2+^ activation cues that are directly modulated by autonomic receptor stimulation. Specifically, βARs and CRs not only, respectively, reduce or increase mean APFI, but also, respectively, shift variability within distributions of Ca^2+^ and V_m_ functions in the same direction (S. Sirenko et al., 2016) as the shift in mean APFI.

### Numerical modeling

Because we experimentally measured characteristics of APs in populations of single cells that differed by autonomic input status, we were able to glean insights not only into APFI variability in an “average” cell, but also into populations of cells isolated from SAN tissue. Embracing SAN function at the cell population level resonates with recent studies of SAN function at the tissue level (Bychkov et al., 2020) (Fenske et al., 2020) that have revealed a novel understanding of the SAN impulse as an emergent property created by a collective of variable interactions among heterogeneous cell populations within the SAN tissue. On the other end, APFI variability per se, also emerges at smaller, subcellular scales, due to variability in the functions of individual molecules, such as ion channels, transporters, and pumps, individually and in complex cooperation with each other. These molecular functions cannot be measured directly in our single cell experiments. Thus, we employed numerical modeling to extend our perspectives from cell populations and single cell levels downwards to the molecular scale. Such broader consideration of variability makes sense when we approach pacemaker function as a multiscale phenomenon (Qu, Garfinkel, Weiss, & Nivala, 2011) featuring free scale and fractal-like characteristics (Weiss & Qu, 2020). Considering the SAN cell as a stochastic dynamical system, we examined variability of major ion currents and submembrane [Ca^2+^] during different autonomic states that created a broad range of AP firing intervals, that were measured experimentally.

Our simulations indicate that the APFI variability of some ion currents and sub-membrane Ca^2+^ can be close to that of the APFI itself, but also can be substantially lower or higher than the APFI variability, depending on the presence or absence of autonomic receptor stimulation, and the time during the AP cycle (Figure 12): components such as I_f_, I_NCX_, I_CaT_ and Ca^2+^ exhibit substantial cycle to cycle variability, whereas I_CaL_, I_KACh_, and I_Kr_ show less, or moderate variability. This behavior reflects complex non-linear recursive interactions of V_m_ and Ca^2+^ that couple the clocks that drive the system (Lyashkov et al., 2018) and, as such, cannot be directly and definitively interpreted in cause-effect terms. Nevertheless, our simulations confirmed ion channel behavior that could have been envisioned. For example, independent of the nature of the noise added (to I_tot_ or to Ca^2+^ release flux): components contributing to DD (I_f_, I_NCX_, and I_CaT_) exhibit larger cycle to cycle variability, whereas components contributing to generation of all-or-none AP characteristics exhibit less variability (peak I_CaL_ and I_Kr_). This result is in accord with the idea of order/disorder transitions during AP cycle (Figure 2, order/disorder dash line), i.e., the DD manifests disorder and transition to order, and hence, a larger variability; then following the AP upstroke, the AP itself manifests order and hence, less variability.

But our simulations also provided some unexpected, interesting results: I_NCX_ and sub-membrane Ca^2+^ amplitudes during DD followed a power law relationship over a wide range of APFI under all conditions tested, indicating that self-similar scale-free or fractal-like behavior is operative within the coupled-clock mechanism via I_NCX_ (Figure 13). There is also likely to be a secondary effect of I_NCX_ amplitude itself on DD acceleration in a recursive fashion. Indeed, an increase in I_NCX_ is expected to accelerate DD, but at the same time, is further accelerated by the very same acceleration it imparts to the ignition mechanism (i.e. diastolic I_CaL_ and Ca^2+^-induced Ca^2+^ release). This self-acceleration ignition mechanism results in the power law behavior predicted by the model.

Peak I_f_ also followed a power function over a broad range of APFIs, albeit in a noisy manner manifesting some extremely long APFIs in CCh. The current that fluctuated most with respect to variabilities in APFI turned out to be I_CaT_, the peak amplitude of which also reflects DD dynamics. When DD is rapidly accelerating (as it does at shorter cycles) I_CaT_ peak quicky activates and achieves a higher peak amplitude; and vice-versa, when DD is slower at longer APFIs I_CaT_ becomes inactivated over time without achieving a peak. In log-log plots this forms a straight-line for almost the entire range of APFIs (power law behavior), except extremely long APFIs when variations in (already) slow DD dynamics have almost no effect on I_CaT_ peak. Thus, simulations of biophysical components within the pacemaker cell system exhibited a power law behavior over a wide range of APFIs that encompasses the broad range of APFIs measured experimentally.

Observation of model simulations through analytic lenses applied to experimental data indicated that (as was the case for experiment data) the simulated variables are self-similar to each other across broad range of APFI within the three autonomic states (Figure 15, Supplementary Figure S4, Supplementary Table S3).

## Model limitations

Our I_CaL_ model was adopted from the Kurata model (Kurata, Hisatome, Imanishi, & Shibamoto, 2002), and while it does include Ca^2+^-dependent inactivation mechanism, it lacks I_CaL_ facilitation described in 2000 by Mangoni et al.(Mangoni et al., 2000). Also, our numerical model of I_f_ lacks dynamic regulation by cAMP (DiFrancesco, 1999), Ca^2+^ activated K^+^ channels and store-operated channels that may potentially contribute to APFIV.

## Author Contributions

EGL and VAM designed the project; DY, AEL, IZ, YY, TMV and BDZ performed experiments and analyzed the results; DY, CHM and ST did the statistical analyses; DY and VAM performed the numerical simulation; DY and EGL wrote the manuscript with the support from YY and VAM. All authors contributed to the article and approved the submitted version.

## Abbreviations

AP: action potential
APD_90_: action potential duration from AP overshoot to 90% repolarization
APFI: AP firing interval
APFIV: variability of AP firing intervals
TTIO: time from the previous AP overshoot to the ignition onset when dV/dt=0.15 (V/S)
CaTFI: firing internal of AP-induced Ca^2+^ transient
CaT_90_: 90% decay time of AP-induced Ca^2+^ transient
LCR: local Ca^2+^ releases
SR: sarcoplasmic reticulum
SD: standard deviation
CV: coefficient of variation
PC: principal component
I_CaL_: high voltage-activated, L-type Ca^2+^ current
I_f_: hyperpolarization-activated funny current
I_KACh_: acetylcholine-activated K^+^ current
I_Kr_: K^+^ current exhibiting strong inward rectification
I_NCX_: Na^+^-Ca^2+^ exchanger (NCX) current
βARs: β-adrenergic receptor stimulation
CRs: Cholinergic receptor stimulation
CCh: carbachol
ISO: isoproterenol
HR: heart rate
SAN: sinoatrial node

## Acknowledgements

The authors wish to acknowledge the assistance of Jia-Hua Qu, MD, PhD for assistance with the PC analysis illustrations, and sincerely appreciate Loretta Lakatta, R.N., B.S.N., Robert Monticone, B.S., Tracy Oppel, and Ruth Sadler, B.A. for their editorial assistance. This manuscript has been released as a pre-print at bioRxiv (https://www.biorxiv.org/content/10.1101/2020.09.01.277756v1) (D. Yang et al., 2020).

