## supplement file for "Ca^2+^ and Membrane Potential Transitions During Action Potentials are Self-Similar to Each Other and to Variability of AP Firing Intervals Across the Broad Physiologic Range of AP Intervals During Autonomic Receptor Stimulation"

**Supplementary Material**

**Supplementary Materials and Methods**

*Single, isolated rabbit sinoatrial nodal cells (SANC) Isolation.* Single, spindle-shaped, spontaneously beating SANC were isolated from the hearts of New Zealand rabbits (Charles River Laboratories, Wilmington, MA) as described previously (Vinogradova et al., 2008). New Zealand White rabbits weighing 2.8~3.1kg were deeply anesthetized with sodium pentobarbital (50~90 mg/kg). The heart was removed quickly and placed in Tyrode solution containing (in mmol/L): NaCl, 130; NaHCO_3_, 24; NaH_2_PO_4_, 1.2; MgCl_2_, 1.0; CaCl_2_, 1.8; KCl, 4.0; and glucose, 5.6, which was saturated with a mixture of 95% 0_2_ and 5% C0_2_ (pH=7.4 at 36^o^C). The sinoatrial node region was cut into ~1.0 mm-wide strips, perpendicular to the crista terminalis. The sinoatrial node preparation was washed twice in Ca^2+^-free Tyrode solution (34^o^C) containing (in mmol/L): NaCl, 140; KCl, 5.4; MgCl_2_, 0.5; NaH_2_PO4, 0.33; HEPES, 5; glucose, 5.5 (pH=6.9); and then incubated at 34^o^C for 30 min in Ca^2+^-free Tyrode solution containing elastase type IV (0.6 mg/ml; Sigma, Chemical Co.), collagenase type 2 (0.8mg/ml; Worthington, NJ, USA), protease type XIV (0.18 mg/ml, Sigma, Chemical Co.) and 0.1% bovine serum albumin (Sigma, Chemical Co.). Thereafter, the sinoatrial node preparation was washed in modified Kraft-Bruhe (KB) solution containing (in mmol/L): potassium glutamate, 70; KCl, 30; KH_2_PO_4_, 10; MgC1_2_, 1; taurine, 20; glucose, 10; EGTA, 0.3 and HEPES, 10 (pH=7.4 with KOH), and kept at 4^o^C for 1h in KB solution containing 50 mg/ml polyvinylpyrrolidone (PVP 40, Sigma, Chemical Co.). Cells were dispersed from the sinoatrial node preparation by gentle pipetting in the KB solution and stored at 4^o^C. Studies in freshly isolated SANC were performed within hours following isolation.

*Spontaneous AP Recordings.* Spontaneous APs (number of beats >512) were recorded in subset-sets of SANC using the perforated patch-clamp technique with an Axopatch 200B patch-clamp amplifier (Axon Instruments) (Bogdanov et al., 2006). The pipette (~3 MΩ) filling solution contained (in mmol/L) K-gluconate 120, NaCl 10, MgATP 5, HEPES 5, KCl 20 (pH, 7.2) and 50 μmol/L β-escin (Sigma). Bath superfusion solution contained (in mmol/L): NaCl 140, KCl 5.4, MgCl_2_ 1; HEPES 5, CaCl_2_, 1.8, Glucose 5.5 (pH=7.4). All functional measurements were performed at 34 ± 0.5^o^C. Both f- and c- SANC were selected for study based on the apparent regularity of their spontaneous beating, the relative smoothness and apparent quality of the cellular surface. Prior to each recording, the spontaneous AP firing was monitored for at least 20 min to insure its stability. The variability of selected AP parameters was measured via a customized program (Bogdanov et al., 2006). In addition to the classic AP characteristics, TTIO (Time to ignition onset) that reflects the onset of the ignition phase of the AP cycle (Lyashkov, Behaer, Lakatta, Yaniv, & Maltsev, 2018) was measured. AP firing interval variability was determined from AP recordings over 20 minutes AP beating time series. AP firing interval variability was characterized by the standard deviations (SD) or coefficient of variation (CV, the ratio of SD to the mean).

*Ca^2+^ Measurements.* Subsets SANC were loaded with the Ca^2+^ indicator fluo-4/AM (5 μmol/L, 20min at room temperature, Thermo Scientific) (Vinogradova et al., 2004). Following washout of extracellular fluo-4/AM, AP initiated global Ca^2+^ transients and spontaneous local Ca^2+^ releases (LCR) during diastole were measured at 34±0.5^o^C with a confocal microscope (Zeiss LSM510, Germany) in the line-scan mode, with the scan line oriented along the long axis of the cell, close to the sarcolemma membrane. Ca^2+^-Image processing, data analysis, and presentation were performed using our original programs written in IDL 6.1 software (Vinogradova et al., 2010) (Interactive Data Language, Harris Corporation). The interval between the peaks of two adjacent AP-triggered Ca^2+^-transients is defined as Ca^2+^-transient firing interval (CaTFI). The LCR period is defined as the time from the peak of the prior AP-induced Ca^2+^ transient to an LCR peak in diastole.

*Drugs.* The synthetic, non-specific β-adrenergic receptor agonist isoproterenol (ISO), cholinergic agonist carbachol (CCh) were from SIGMA.

**Supplementary Discussion**

***Links between clock membrane potential and Ca^2+^ functional parameters measured during spontaneous AP firing and their molecular underpinnings.*** The rapid depolarization from the diastolic membrane potential during the AP upstroke, mainly informs on the availability of Ca_V_1.2 L-type Ca^2+^ channels for activation by the acute depolarization (Adeniran, McPate, Witchel, Hancox, & Zhang, 2011; DiPolo & Beauge, 2006; Faber, Silva, Livshitz, & Rudy, 2007) (Fig. S2). The greater the availability of channels to respond to a membrane voltage change cue, i.e. to than can be synchronously activated by a membrane voltage change (greater Ca^2+^ channel availability), the greater **dV/dt** of AP upstroke. The resultant Ca^2+^ influx via activated L-type Ca^2+^ channels binds to RyRs and synchronizes their activation via Ca^2+^-induced Ca^2+^ release (CICR), resulting in graded, synchronous RyR Ca^2+^ release that generates the AP-triggered **CaT** (Koivumaki, Korhonen, Takalo, Weckstrom, & Tavi, 2009; Stern et al., 1999) (Fig. S2), depleting the Ca^2+^ charge on the SR Ca^2+^ capacitator. L-Type channels also begin to inactivate with time, even at the depolarized membrane potential, facilitated by the synchronous RyR Ca^2+^ release via CICR. The AP-induced cytosolic transient [Ca^2+^] due to RyR Ca^2+^ release and Ca^2+^ influx via L-type Ca channels activates Ca^2+^ ATPase (Serca) to pump Ca^2+^ into SR, and the kinetics of pumping that recharge the Ca^2+^ capacitor, as reflected in **CaT_90_** (Fig. S2). Because **CaT_90_** of the AP-induced Ca^2+^ transient informs, in part, on the kinetics of SR Ca^2+^ refilling following Ca^2+^ depletion by the prior AP (Vinogradova et al., 2010), gradations in the mean CaT_90_ of a given “steady state” are also likely linked to gradations of beat-to-beat Ca-ATPase availability to become activated by Ca^2+^ (Fig. S2). Thus, beat-to-beat variability of Ca^2+^ removal from the cytosol and pumping into SR is one mechanism that may lead to variability of local LCR periods (Fig. S2).

Surface membrane depolarization activates K^+^ channels and inactivates forward Na-Ca exchange: AP repolarization kinetics, e.g. **APD_90_**, inform on the combined actions of K^+^ channels, cytosolic [Ca^2+^], I_NCX_ and I_f_ (Fig. S2). K^+^ channel activation repolarizes the surface membrane, reactivating forward Na-Ca exchange, which assists in removing the Ca^2+^ released via the AP-induced RyR flux (which, in a “steady state” is equal to the amount of all Ca^2+^ influx via L-Type Ca^2+^ channels). Ca^2+^ also activates K^+^ channels, assisting in cell membrane repolarization (**APD_90_**), even prior to achieving the full surface membrane repolarization (i.e. MDP) and full I_f_ activation. Time from AP upstroke to MDP is regulated by the aforementioned ensemble of molecular activation and inactivation.

Following the reestablishment of the MDP, the membrane potential begins to slowly depolarize (Fig. S2), due to removal of K^+^ channel activation, I_f_ activation, and spontaneous asynchronous local RyR activation that generates submembrane LCRs (Fig. S2). **LCR periods** inform not only on the RyR activation but also on the kinetics of recharging the SR Ca^2+^ capacitor (Vinogradova et al., 2010). Summation of individual LCRs regulated, in part, by CICR (Stern et al., 2014), produces an LCR ensemble Ca^2+^ signal that activates an inward I_NCX_. T-type (Ca_V_3.1) and low voltage-activated L-Type channels (Ca_V_1.3) become activated during diastolic depolarization and likely contribute to the increase in submembrane [Ca^2+^]. Exponential growth of the LCR ensemble Ca^2+^ signal (Fig. S2), exponentially increasing I_NCX_ activation, initiating a rapid acceleration of the DD rate (non-linear DD component) (Lyashkov et al., 2018). Thus, **TTIO** informs on the kinetics of LCR generation and synchronization to generate the LCR ensemble Ca^2+^ signal that increase I_NCX_ activation. I_f_ inactivates during late DD Rapid and concurrent dynamic, non-linear feed-forward interactions of membrane potential, I_f_, Ca^2+^ channel activation and the increase in I_NCX_ due in large measure to exponential growth of the LCR ensemble Ca^2+^ signal, drive the late DD membrane potential to the threshold potential required for activation of Ca_V_1.2 L-Type Ca^2+^ channels and the next rapid AP upstroke ensues (Lyashkov et al., 2018).

**Supplementary Figures**

**Supplementary Figure S1.** Schematic of Experimental Data and Data Analyses

**Supplementary Figure S2. *Molecular drivers of coupled oscillator pacemaker functions.*** The activation-inactivation schema of molecular functions that underlie AP (Adeniran et al., 2011; Altomare et al., 2001; DiPolo & Beauge, 2006; Faber et al., 2007) and Ca^2+^ loop (Koivumaki et al., 2009; Stern et al., 1999) functions.

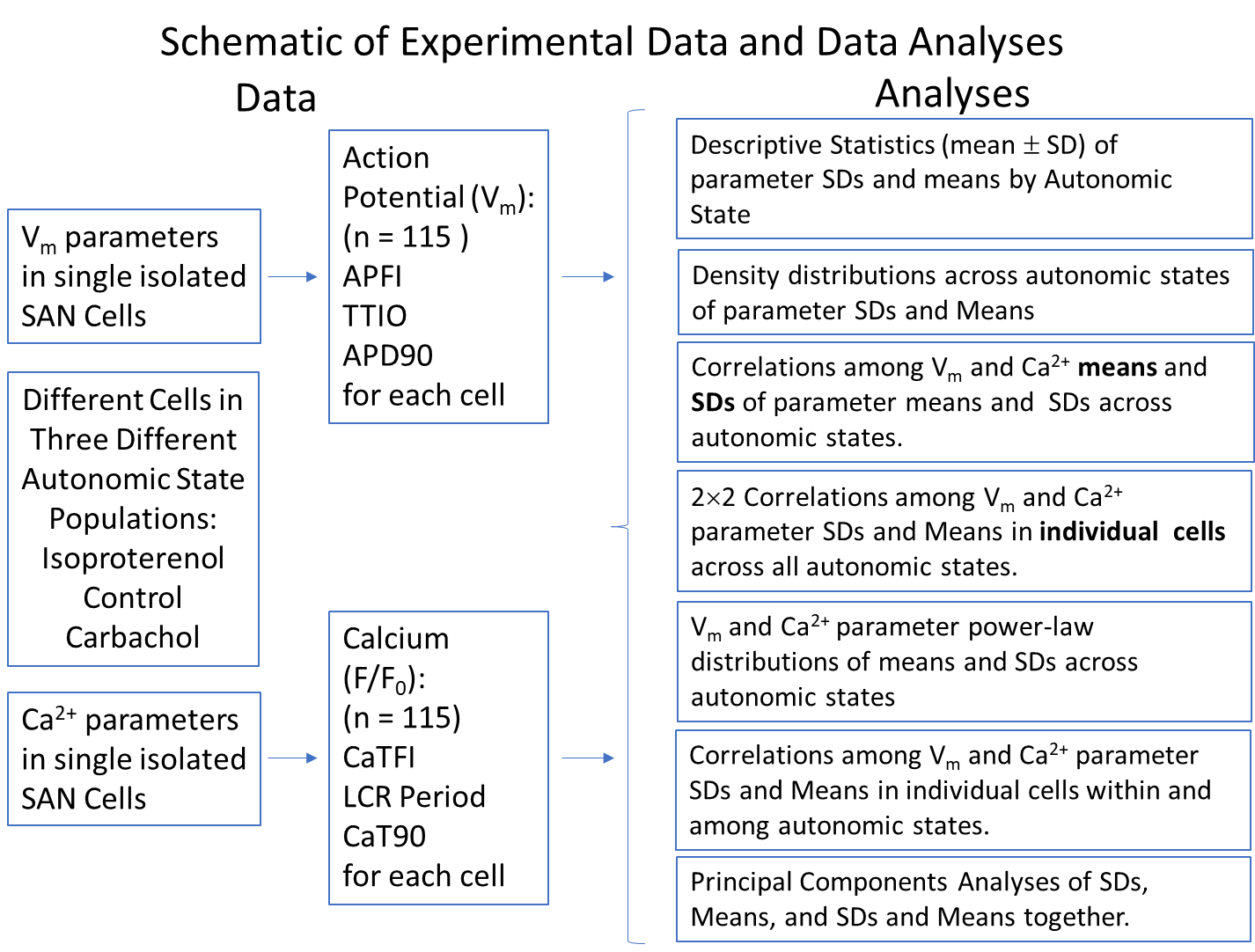

Surface membrane ion channels, electrogenic ion exchange proteins, e.g., NCX, and ion pumps, e.g., Na-K ATPase are ion current oscillators that regulates TTIO (Sirenko et al., 2016 Lyashkov et al., 2018) and APD_90_. The NCX and Na-K ATPase are voltage- and transmembrane gradient-dependent, and actively and exquisitely responsive to effect changes in Na^+^-Ca^2+^ electro-chemical gradient oscillations that underline each AP cycle in isolated SANC (Sirenko et al., 2016). Molecules within the bi-directionally coupled oscillator system either directly or indirectly regulate both surface membrane potential and intracellular Ca^2+^. Variable degrees of self-organized coherence or synchronization among these molecular functions impact on the fidelity to which the Ca^2+^ oscillators within the system and coupled to the system current oscillators (Yaniv, Lyashkov, & Lakatta, 2013; Yaniv et al., 2014).

The SR oscillates intracellular Ca^2+^, that includes the LCR (indicated as the white arrows on Ca^2+^ line-scan image) period and CaT_90_: SR operates as a Ca^2+^ capacitor, its Ca^2+^ charge is regulated by Serca that pumps Ca^2+^ into the SR lumen, and by ryanodine receptors (RyR) that dissipate the Ca^2+^ charge, via releasing Ca^2+^ beneath the cell surface membrane. An interaction of PLB with Serca modulates the maximum speed of Ca^2+^ pumping into SR. Ion pumps that operate within each clock are energy dependent. The LCR period is an index of clock coupling that becomes manifest when a SANC fires an AP.

**Supplementary Figure S3.** Heatmaps of correlations by group.

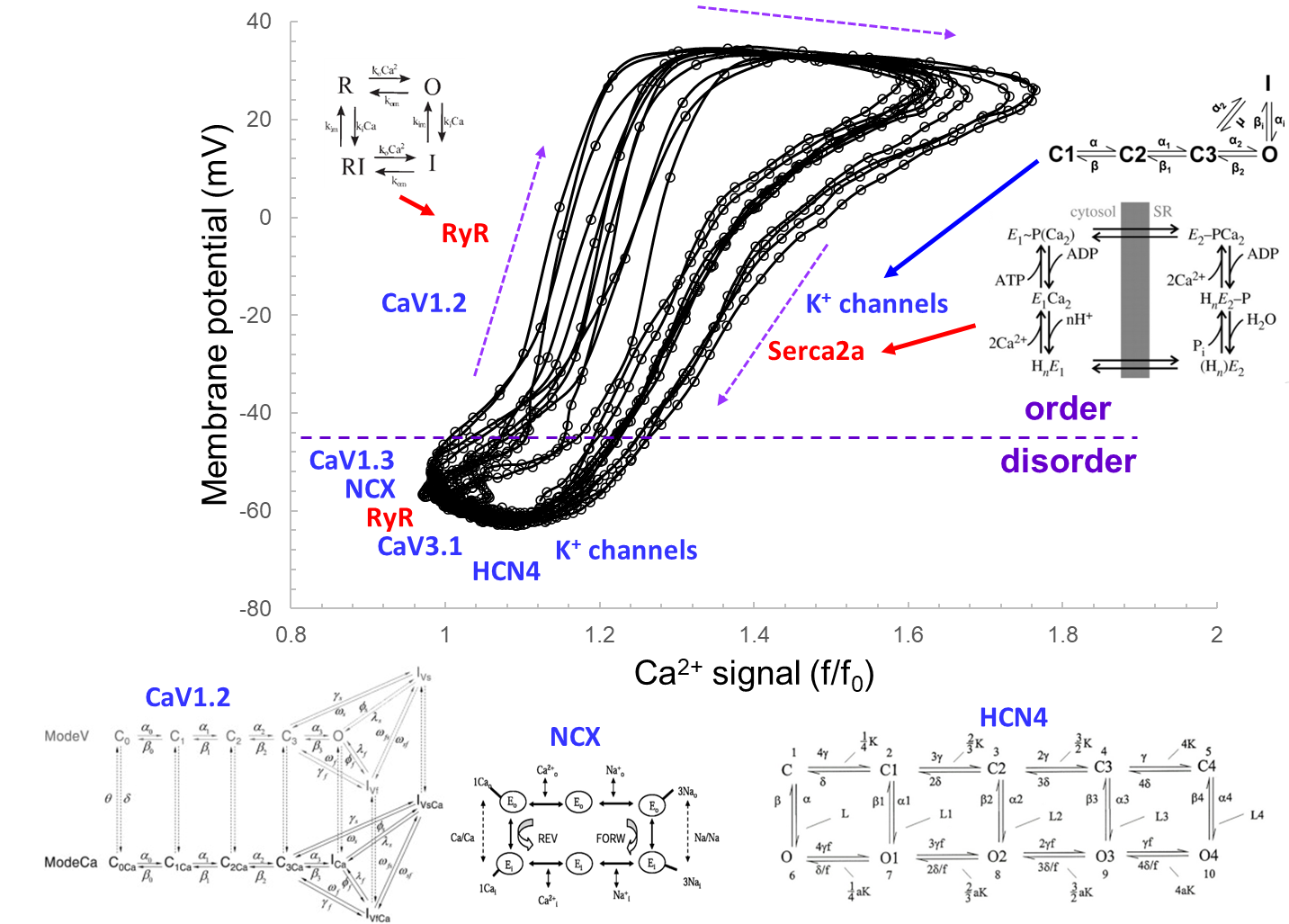

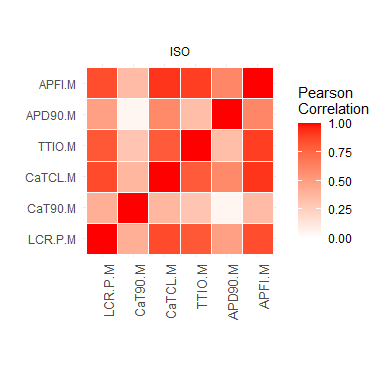

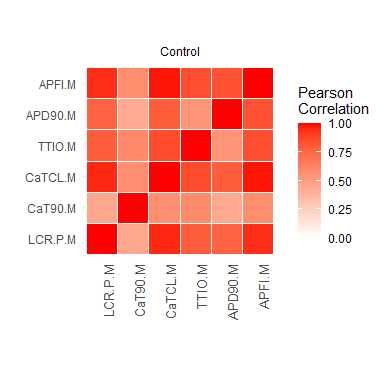

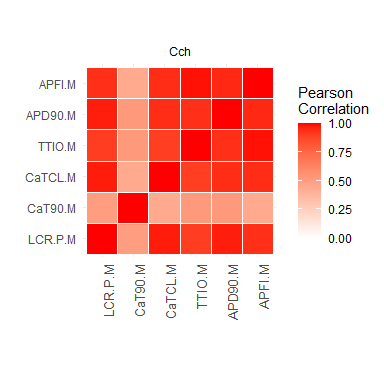

**Supplementary Figure S4.** PCA of modeling data: only 2 PCs explain 93.9% of the variation in these 8 variables!

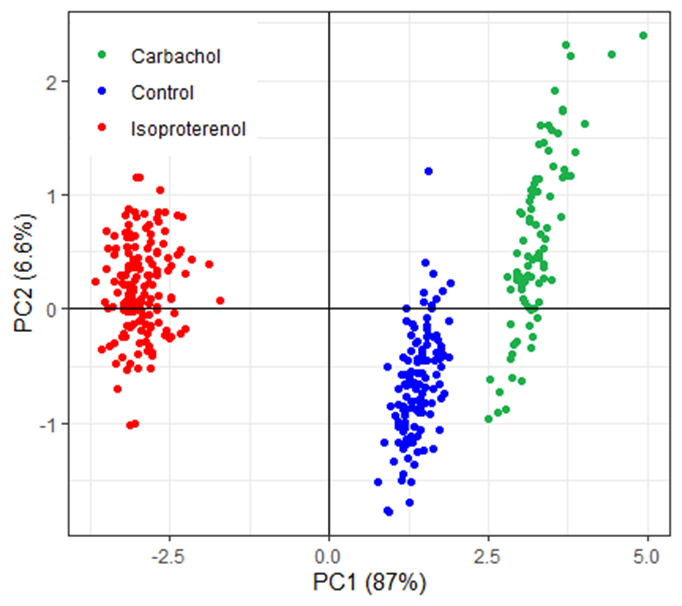

**Supplementary Figure S5.** A Comparison of gradations in the phosphorylation level (**A**) indexed by phosphorylated PLB at Ser^16^ normalized to total PLB of cells in different steady states (redrawn from published previously data, (Lyashkov et al., 2009; Yang et al., 2012)) and the means (**B**) and SDs (**C**) of all 6 parameters in all 3 “steady states” listed in Table 1. Phosphorylation data and functional data are normalized to value in control, as indicated by the dashed blue line in figure panels.

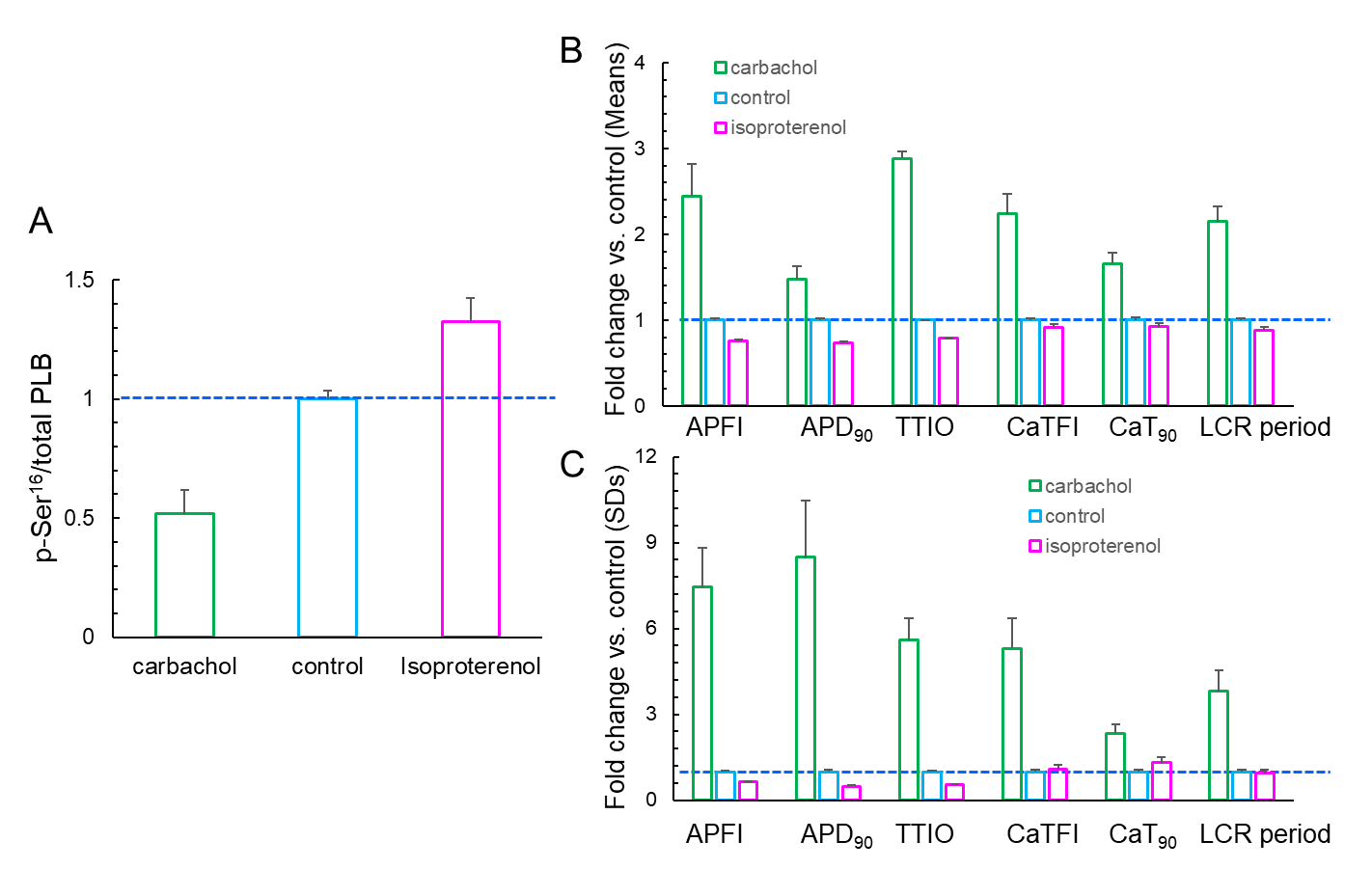

**Supplementary Table S1**. **Supplement table S1.** Sd1 and sd2 from Ellipse fitting of Poincaré Plots from AP recordings during CCh or ISO super-perfusion and its self-control (the same cells as Figures 4-6).

|  |  | **sd1** | **sd2** | **sd1/sd2** | **Mean** | **SD** |
| --- | --- | --- | --- | --- | --- | --- |
| **APFI** | **CCh** | 39.922 | 60.755 | 0.6571 | 910.279 | 53.146 |
| (ms) | **Con for CCh** | 13.501 | 15.381 | 0.8778 | 330.378 | 14.455 |
|  | **Con for ISO** | 7.270 | 13.605 | 0.5344 | 305.486 | 10.906 |
|  | **ISO** | 2.868 | 6.091 | 0.4708 | 192.680 | 4.758 |
| **APD90** | **CCh** | 27.061 | 30.152 | 0.8975 | 269.898 | 28.595 |
| (ms) | **Con for CCh** | 8.037 | 8.509 | 0.9445 | 161.196 | 8.266 |
|  | **Con for ISO** | 5.124 | 6.159 | 0.8320 | 142.428 | 5.661 |
|  | **ISO** | 1.815 | 4.226 | 0.4295 | 106.104 | 3.250 |
| **TTIO** | **CCh** | 69.653 | 105.826 | 0.6582 | 848.680 | 90.217 |
| (ms) | **Con for CCh** | 16.732 | 28.683 | 0.5833 | 258.300 | 24.099 |
|  | **Con for ISO** | 14.689 | 23.947 | 0.6134 | 242.383 | 19.875 |
|  | **ISO** | 5.678 | 8.342 | 0.6807 | 145.631 | 7.132 |

**Supplementary table S2:** Summary Table for Principal Component Analyses.

| Data | % Explained,  Cumulative % | SDs | Means | Means & SDs |
| --- | --- | --- | --- | --- |
| **Vm data** | PC1 | 89.9 | 91.7 | 85.9 |
|  | PC2 | 8.0, 98.0 | 7.9, 99.6 | 6.8, 92.7 |
|  | PC3 | 2.0, 100.0 | 0.4, 100.0 | 4.4, 97.1 |
| **Ca^2+^ data** | PC1 | 72.0 | 86.4 | 70.7 |
|  | PC2 | 18.2, 90.2 | 12.9, 99.3 | 11.9, 82.6 |
|  | PC3 | 9.8, 100.0 | 0.7, 100.0 | 9.2, 91.9 |
| **Vm and Ca^2+^ data** | PC1 | 68.4 | 85.8 | 76.7 |
|  | PC2 | 16.1, 84.6 | 7.8, 93.6 | 9.5, 83.2 |
|  | PC3 | 6.9, 91.5 | 4.1, 97.8 | 5.7, 88.8 |

**Supplementary table S3:** Correlations of modeling data

| Correlations and p-values | APFI | TTIO | INCX.m40 | If.m40 | IKr.m40 | ICaL.m40 | ICaT.m40 | Ca.m40.mM |
| --- | --- | --- | --- | --- | --- | --- | --- | --- |
| APFI : r | 1 | 0.7269 | 0.8556 | 0.8777 | -0.8324 | 0.6578 | 0.8365 | -0.8488 |
| APFI : p | NA | 0 | 0 | 0 | 0 | 0 | 0 | 0 |
| TTIO : r | 0.7269 | 1 | 0.6951 | 0.713 | -0.7081 | 0.5833 | 0.6618 | -0.6905 |
| TTIO : p | 0 | NA | 0 | 0 | 0 | 0 | 0 | 0 |
| INCX.m40 : r | 0.8556 | 0.6951 | 1 | 0.9387 | -0.9789 | 0.8895 | 0.987 | -0.9999 |
| INCX.m40 : p | 0 | 0 | NA | 0 | 0 | 0 | 0 | 0 |
| If.m40 : r | 0.8777 | 0.713 | 0.9387 | 1 | -0.9502 | 0.8639 | 0.9225 | -0.9346 |
| If.m40 : p | 0 | 0 | 0 | NA | 0 | 0 | 0 | 0 |
| IKr.m40 : r | -0.8324 | -0.7081 | -0.9789 | -0.9502 | 1 | -0.9228 | -0.9521 | 0.9778 |
| IKr.m40 : p | 0 | 0 | 0 | 0 | NA | 0 | 0 | 0 |
| ICaL.m40 : r | 0.6578 | 0.5833 | 0.8895 | 0.8639 | -0.9228 | 1 | 0.8621 | -0.8909 |
| ICaL.m40 : p | 0 | 0 | 0 | 0 | 0 | NA | 0 | 0 |
| ICaT.m40 : r | 0.8365 | 0.6618 | 0.987 | 0.9225 | -0.9521 | 0.8621 | 1 | -0.9873 |
| ICaT.m40 : p | 0 | 0 | 0 | 0 | 0 | 0 | NA | 0 |
| Ca.m40.mM : r | -0.8488 | -0.6905 | -0.9999 | -0.9346 | 0.9778 | -0.8909 | -0.9873 | 1 |
| Ca.m40.mM : p | 0 | 0 | 0 | 0 | 0 | 0 | 0 | NA |

**DESCRIPTION OF NUMERICAL MODEL**

We performed simulations using modified Maltsev-Lakatta SA node cell numerical model that features a coupled-clock mechanism (Maltsev and Lakatta, 2009). The original model is freely available and can be downloaded and run in CellML format (<http://models.cellml.org/workspace/maltsev_2009>) using the Cellular Open Resource software developed by Alan Garny at Oxford University in the UK (Garny et al., 2009) (for recent development of this software see <http://www.opencor.ws/>). We simulated APs, ion currents, and Ca dynamics in three scenarios: basal AP firing, β adrenergic receptor (βAR) stimulation with isoproterenol (ISO), 100 nM, and cholinergic receptor (CR) stimulation with carbachol (CCh), 0.1 μM. The effect of ISO was modelled as previously described in our previous studies (Maltsev and Lakatta, 2010), but the effect of CR stimulation on I_CaL_ was changed to Zaza et al. model (Zaza et al., 1996) as described in details below. All model equations and parameter values are provided below.

**MODEL PARAMETERS**

**Fixed ion concentrations, mM**

*Ca*_o_ = 2: Extracellular Ca^2+^ concentration.

*K*_o_ = 5.4: Extracellular K^+^ concentration.

*K*_i_=140: Intracellular K^+^ concentration.

*Na*_o_ = 140: Extracellular Na^+^ concentration.

*Na*_i_=10: Intracellular Na^+^ concentration.

*Mg*_i_ = 2.5: Intracellular Mg^2+^ concentration.

**Cell compartments**

*C*_m_ = 32 pF: Cell electric capacitance.

*L*_cell_ = 70 μm: Cell length.

*R*_cell_ = 4 μm: Cell radius.

*L*_sub_ = 0.02 μm: Distance between jSR and surface membrane (submembrane space).

*V*_cell_ = π·*R*_cell_^2^·*L*_cell_ = 3.5185838 pL: Cell volume.

*V*_sub_ = 2π·*L*_sub_·(*R*_cell_ - *L*_sub_/2)·*L*_cell_ = 0.035097874 pL: Submembrane space volume.

*V*_jSR_part_ = 0.0012: Part of cell volume occupied by junctional SR.

*V*_jSR_ = *V*_jSR_part_·*V*_cell_: Volume of junctional SR (Ca^2+^ release store).

*V*_i_part_ = 0.46: Part of cell volume occupied with myoplasm.

*V*_i_ = *V*_i_part_·*V*_cell_-*V*_sub_: Myoplasmic volume.

*V*_nSR_part_ = 0.0116: Part of cell volume occupied by network SR.

*V*_nSR_ = *V*_nSR_part_·*V*_cell_: Volume of network SR (Ca^2+^ uptake store).

**The Nernst equation and electric potentials, mV**

E_X_ = (RT/F) · ln([X]_o_/[X]_i_) = *E*_T_ · ln([X]_o_/[X]_i_), where

F = 96485 C/M is Faraday constant,

T = 310.15 K˚ is absolute temperature for 37˚C,

R = 8.3144 J/(M·K˚) is the universal gas constant,

*E*_T_ is “RT/F” factor = 26.72655 mV,

and [X]_o_ and [X]_i_ are concentrations of an ion “X” out and inside cell, respectively.

*E*_Na_ = *E*_T_ ∙ ln(Na_o_/Na_i_): Equilibrium potential for Na^+^.

*E*_K_ = *E*_T_ ∙ ln(K_o_/K_i_): Equilibrium potential for K^+^.

*E*_Ks_ = *E*_T_ ∙ ln[(K_o_ + 0.12∙ Na_o_)/(K_i_ + 0.12 ∙ Na_i_)]: Reversal potential of *I*_Ks_.

*E*_CaL_ = 45: Apparent reversal potential of *I*_CaL_.

*E*_CaT_ = 45: Apparent reversal potential of *I*_CaT_.

*E*_st_ = 37.4: Apparent reversal potential of *I*_st_.

**Sarcolemmal ion current types and their parameter values**

*I*_CaL_:	L-type Ca^2+^ current [*g*_CaL,max,basal_ = 0.58 nS/pF, as in Kurata et al. model (Kurata et al., 2002)].

Steady-state activation parameters: *V*_½,d_ =-13.5 mV; *K*_d_ =6 mV.

Steady-state inactivation parameters: *V*_½,f_ =-35 mV; *K*_f_ =7.3 mV.

*K*_mfCa_ = 0.00035 mM: Dissociation constant of Ca^2+^ -dependent *I*_CaL_ inactivation.

*β*_fCa_ = 60 mM^-1^ · ms^-1^: Ca^2+^ association rate constant for *I*_CaL_.

*α*_fCa_ = 0.021 ms^-1^ : Ca^2+^ dissociation rate constant for *I*_CaL_

*I*_CaT_:	T-type Ca^2+^ current (*g*_CaT,max_ = 0.1832 nS/pF).

*I*_f_:	Hyperpolarization-activated current (*g*_If,max_ = 0.15 nS/pF).

*V*_If,1/2,basal_ = -64 mV: half activation voltage for *I*_f_ current in the basal state.

*s*_max_ = -7.2 mV: maximum ACh-induced shift of *I*_f_ half activation voltage.

*n_f_* = 0.69 and *K_0.5,f_* = 12.6 nM: Michaelis-Menten parameters for ACh modulation of *I*_f_.

*I*_st_:	Sustained non-selective current (*g*_st,max_ = 0.003 nS/pF).

*I*_Kr_:	Delayed rectifier K^+^ current rapid component (*g*_Kr,max_ = 0.08113973 nS/pF).

*I*_Ks_:	Delayed rectifier K^+^ current slow component (*g*_Ks,max_ = 0.0259 nS/pF).

*I*_to_:	4-aminopyridine sensitive transient K^+^ current (*g*_to,max_ = 0.252 nS/pF).

*I*_sus_:	4-aminopyridine sensitive sustained K^+^ current (*g*_sus,max_ = 0.02 nS/pF).

*I*_NaK_:	Na^+^/K^+^ pump current (*I*_NaK,max_ = 2.88 pA/pF).

*K*_mKp_ = 1.4 mM: Half-maximal *K*_o_ for *I*_NaK_.

*K*_mNap_ = 14 mM: Half-maximal *Na*_i_ for *I*_NaK_.

*I*_bCa_:	Background Ca^2+^ current (*g*_bCa_ = 0.0006 nS/pF).

*I*_bNa_:	Background Na^+^ current (*g*_bNa_ = 0.00486 nS/pF).

*I*_KACh_:	Acetylcholine-activated K^+^ current; *I*_KACh_ =0, when [ACh]=0.

*g*_KAch,max_ =0.14241818 nS/pF.

*I*_NCX_:	Na^+^/Ca^2+^ exchanger (NCX) current (*k*_NCX_ = 187.5 pA/pF).

*K*_1ni_ = 395.3: intracellular Na^+^ binding to first site on NCX.

*K*_2ni_ = 2.289: intracellular Na^+^ binding to second site on NCX.

*K*_3ni_ = 26.44: intracellular Na^+^ binding to third site on NCX.

*K*_1no_ = 1628: extracellular Na^+^ binding to first site on NCX.

*K*_2no_ = 561.4: extracellular Na^+^ binding to second site on NCX.

*K*_3no_ = 4.663: extracellular Na^+^ binding to third site on NCX.

*K*_ci_ = 0.0207: intracellular Ca^2+^ binding to NCX transporter.

*K*_co_ = 3.663: extracellular Ca^2+^ binding to NCX transporter.

*K*_cni_ = 26.44: intracellular Na^+^ and Ca^2+^ simultaneous binding to NCX.

*Q*_ci_:= 0.1369: intracellular Ca^2+^ occlusion reaction of NCX.

*Q*_co_=0: extracellular Ca^2+^ occlusion reaction of NCX.

*Q*_n_= 0.4315: Na^+^ occlusion reactions of NCX.

**Ca^2+^ diffusion**

*τ*_difCa_ = 0.04 ms: Time constant of Ca^2+^ diffusion from the submembrane to myoplasm.

*τ*_tr_ = 40 ms: Time constant for Ca^2+^ transfer from the network to junctional SR.

**SR Ca^2+^ ATPase function**

*K*_up_ = 0.6·10^-3^ mM: Half-maximal Ca_i_ for Ca^2+^ uptake in the network SR.

*P*_up,basal_ = 0.012 mM/ms: Rate constant for Ca^2+^ uptake by the Ca^2+^ pump in the network SR.

**RyR function**

*k*_oCa_ = 10 mM^-2^· ms^-1^; *k*_om_ = 0.06 ms^-1^; *k*_iCa_ = 0.5 mM^-1^· ms^-1^ ; *k*_im_ = 0.005 ms^-1^; *EC*_50_SR_ = 0.45 mM; *k*_s_ = 250·10^3^ ms^-1^; *MaxSR =*15; *MinSR =*1; *HSR =* 2.5;

**Ca^2+^ and Mg^2+^ buffering**

*k*_bCM_=0.542 ms^-1^: Ca^2+^ dissociation constant for calmodulin.

*k*_bCQ_=0.445 ms^-1^: Ca^2+^ dissociation constant for calsequestrin.

*k*_bTC_=0.446 ms^-1^: Ca^2+^ dissociation constant for the troponin-Ca^2+^ site.

*k*_bTMC_=0.00751 ms^-1^: Ca^2+^ dissociation constant for the troponin-Mg^2+^ site.

*k*_bTMM_=0.751 ms^-1^: Mg^2+^ dissociation constant for the troponin-Mg^2+^ site.

*k*_fCM_=227.7 mM^-1^· ms^-1^: Ca^2+^ association constant for calmodulin.

*k*_fCQ_=0.534 mM^-1^· ms^-1^: Ca^2+^ association constant for calsequestrin.

*k*_fTC_=88.8 mM/ms: Ca^2+^ association constant for troponin.

*k*_fTMC_=227.7 mM/ms: Ca^2+^ association constant for the troponin-Mg^2+^ site.

*k*_fTMM_=2.277 mM/ms: Mg^2+^ association constant for the troponin-Mg^2+^ site.

*TC*_tot_=0.031 mM: Total concentration of the troponin-Ca^2+^ site.

*TMC*_tot_=0.062 mM: Total concentration of the troponin-Mg^2+^ site.

*CQ*_tot_=10 mM: Total calsequestrin concentration.

*CM*_tot_=0.045 mM: Total calmodulin concentration.

**FORMULATIONS: ELECTROPHYSIOLOGY**

**Membrane potential, *V*_m_ (variable *y_15_*)**

**dV_m_*/*dt = -* (*I*_CaL_ + *I*_CaT_ + *I*_f_ + *I*_st_ + *I*_Kr_ + *I*_Ks_ + *I*_to_ + *I*_sus_ + *I*_NaK_ + *I*_NCX_ + *I*_bCa_ + *I*_bNa_+ *I*_KACh_) /*C*_m_

**Gating variables *(y_16_ - y_30_)* and their differential equations**

*dy_i_/dt = (y_i,∞_ -* *y)*/*τ_yi_*

(*y_i_ =* *d*_L_, *f*_L_, *f*_Ca_, *d*_T_, *f*_T_, *p*_aF_, *p*_aS_, *p*_i_, *n*, *q*, *r*, *y*, *q*_a_, *q*_i_, *a*)

*τ_yi_*: Time constant for a gating variable *y_i_*.

*α_yi_* and *β_yi_*: Opening and closing rates for channel gating.

*y_i_,*_∞_: Steady-state curve for a gating variable *y_i_*.

**Ion currents**

**L-type Ca^2+^ current (*I*_CaL_),** based on formulations of Kurata et al. (Kurata et al., 2002) that include Ca^2+^ dependent *I*_CaL_ inactivation. See also Table S4 for comparison of steady-state activation parameters with those in other SANC models.

*I*_CaL_=*C*_m_·*g*_CaL,max_ ·(*V_m_- E*_CaL_)·*d*_L_·*f*_L_·*f*_Ca_

*d*_L,∞_ =1/{1+ exp[-(*V_m_*- *V*_½,d_ )/ *K*_d_ ]}

*f*_L,∞_ =1/{1+exp[(*V_m_* - *V*_½,f_ )/ *K*_f_ ]}

*α*_dL_ = -0.02839·(*V_m_*+ 35)/ {exp[-(*V_m_*+35)/2.5] - 1} -0.0849 · *V_m_* / [exp(-*V_m_*/4.8)- 1]

*β*_dL_ = 0.01143 · (*V_m_* - 5)/ {exp[(*V_m_*- 5)/2.5] -1}

*τ*_dL_ =1/(*α*_dL_ + *β*_dL_)

*τ*_fL_ = 257.1 · exp{-[(*V_m_*+ 32.5)/13.9]^2^ }+ 44.3

*f*_Ca,∞_ =*K*_mfCa_ / (*K*_mfCa_ + *Ca*_sub_)

*τ*_fCa_ = *f*_Ca,∞_ / *α*_fCa_

**βAR stimulation**: I*_CaL_* traces and their respective IV curves in control vs. in the presence of βAR stimulation are shown in Figure S6.

**CR stimulation:** The fractional block (*b*_CaL_) of *I*_CaL_ by CR stimulation was adopted from Zaza et al. model given in Figure 2 legend in (Zaza et al., 1996):

*b*_CaL_ = [ACh]^0.348/(2921^0.348+[ACh]^0.348), where [ACh] is given in μM

*g*_CaL,max_ =*C*_m_·*g*_CaL,basal_ · (1 - *b*_CaL_)

Thus, the fractional block in the presence of 0.1 μM of CCh (simulated in the present study) is expected relatively small: *b*_CaL_(0.1) = 0.02728, i.e. <3% (see Figure S7)

**T-type Ca^2+^ current (*I*_CaT_),** based on formulations suggested by Demir et al., (Demir et al., 1994) and modified by Kurata et al. (Kurata et al., 2002).

*I*_CaT_ = *C*_m_∙*g*_CaT,max_ ∙(*V_m_*- *E*_CaT_)∙*d*_T_∙*f*_T_

*d*_T,∞_ =1/ {1 + exp[-(*V_m_* + 26.3)/6.0]}

*f*_T,∞_ = 1/{1+ exp[(*V_m_*+ 61.7)/5.6]}

*τ*_dT_ = 1/{1.068∙exp[(*V_m_*+ 26.3)/30] + 1.068∙exp[-(*V_m_* + 26.3)/30]}

*τ*_fT_ =1/{0.0153∙exp[- (*V_m_* + 61.7)/83.3] + 0.015∙exp[(*V_m_*+ 61.7)/15.38]}

**Rapidly activating delayed rectifier K^+^ current (*I*_Kr_)**, based on formulations suggested by Zhang et al. (Zhang et al., 2000) and modified by Kurata et al. (Kurata et al., 2002).

*I*_Kr_ = *C*_m_∙*g*_Kr,max_ ∙(*V_m_* - *E*_K_)∙(0.6∙ *p*_aF_ + 0.4∙ *p*_aS_) ∙ *p*_i_

*p*_a,∞_ =1/ {1 + exp[-(*V_m_*+23.2)/10.6]}

*p*_i,∞_ = 1/ {1 + exp[(*V_m_* + 28.6)/17.1]}

*τ*_paF_ = 0.84655354/[0.0372 ∙ exp(*V_m_*/15.9) + 0.00096 ∙ exp(-*V_m_* /22.5)]

*τ*_paS_ = 0.84655354/[0.0042 ∙ exp(*V_m_* /17.0) + 0.00015 ∙ exp(-*V_m_* /21.6)]

*τ*_pi_ = 1/[0.1 ∙ exp(-*V_m_*/54.645) + 0.656 ∙ exp(*V_m_*/106.157)]

**Slowly activating delayed rectifier K^+^ current (*I*_Ks_)**, based on formulations suggested by Zhang et al. (Zhang et al., 2000).

*I*_Ks_ = *C*_m_∙*g*_Ks,max_ ∙(*V_m_* - *E*_Ks_)∙ *n*^2^

*α*_n_ = 0.014/ {1 + exp[-(*V_m_*- 40)/9]}

*β*_n_ = 0.001 ∙ exp(-*V_m_*/45)

*n*_∞_ = *α*_n_/(*α*_n_ + *β*_n_)

*τ*_n_ =1/(*α*_n_ + *β*_n_)

**4-aminopyridine-sensitive currents (*I*_4AP_ =*I*_to_ *+ I*_sus_),** based on formulations suggested by Zhang et al. (Zhang et al., 2000).

*I*_to_ = *C*_m_ ∙ *g*_to,max_ ∙ (*V_m_*- *E*_K_) ∙ *q*∙ *r*

*I*_sus_ = *C*_m_ ∙ *g*_sus,max_ ∙ (*V_m_* - *E*_K_) ∙ *r*

*q*_∞_ =1/{1 + exp[(*V_m_*+ 49)/13]}

*r*_∞_ =1/{1 + exp[-(*V_m_* - 19.3)/15]}

*τ*_q_ = 39.102/{0.57∙exp[-0.08∙(*V_m_*+44)]+0.065∙exp[0.1∙(*V_m_*+45.93)]}+ 6.06

*τ*_r_ =14.40516/{1.037∙exp[0.09∙(*V_m_*+30.61)]+0.369∙exp[-0.12∙(*V_m_*+23.84)]}+ 2.75352

**Hyperpolarization-activated, “funny” current (*I*_f_)**, based on formulations of Wilders at al. (Wilders et al., 1991) and Kurata et al.(Kurata et al., 2002). The shift *s* (in mV) of the *I*_f_ activation curve by ChR stimulation was adopted from Zhang et al. 2002 (Zhang et al., 2002).

*I*_f_ = *I*_fNa_+ *I*_fK_

*y*_∞_ = 1/{1 + exp[(*V_m_* - *V*_If,1/2_) /13.5]}

τ_y_ = 0.7166529/{exp[-(*V_m_*+ 386.9)/45.302] + exp[(*V_m_* - 73.08)/19.231]}

*I*_fNa_ = *C*_m_∙0.3833 ∙*g*_If,max_ ∙(*V_m_* - *E*_Na_)∙*y*^2^

*I*_fK_ = *C*_m_∙0.6167 ∙ *g*_If,max_ ∙(*V_m_* - *E*_K_)∙*y*^2^

*V*_If,1/2_ = *V*_If,1/2,basal_ + *s*

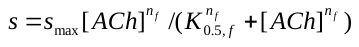

**Sustained inward current (*I*_st_),** based on formulations of Shinigawa et al. (Shinagawa et al., 2000) which were adopted for rabbit SANC by Kurata et al. (Kurata et al., 2002).

*I*_st_ = *C*_m_ ∙*g*_st,max_ ∙ (*V_m_* - *E*_st_)∙ *q*_a_∙ *q*_i_

*q*_a,∞_ =1/{1 + exp[-(*V_m_* + 57)/5]}

*α*_qa_ =1/{0.15 ∙ exp(-*V_m_*/11) + 0.2 ∙ exp(-*V_m_*/700)}

*β*_qa_ =1/{16 ∙ exp(*V_m_*/8) + 15 ∙ exp(*V_m_*/50)}

τ_qa_ =1/(*α*_qa_ + *β*_qa_)

*α*_qi_ =1/{3100 ∙ exp(*V_m_*/13) + 700 ∙exp(*V_m_*/70)}

*β*_qi_ =1/{95∙ exp(-*V_m_*/10) + 50 ∙exp(-*V_m_*/700)} + 0.000229/[1 + exp(-*V_m_*/5)]

*τ*_qi_ =6.65/(*α*_qi_ + *β*_qi_)

*q*_i,∞_ =*α*_qi_ /(*α*_qi_ + *β*_qi_)

**Na^+^-dependent background current (*I*_bNa_)**

*I*_b,Na_ =*C*_m_∙*g*_bNa_∙(*V_m_* - *E*_Na_)

**Na^+^-K^+^ pump current (*I*_NaK_),** based on formulations on Kurata et al. (Kurata et al., 2002), which were in turn based on the experimental work of Sakai et al. (Sakai et al., 1996) for rabbit SANC.

*I*_NaK_ = *C*_m_ ∙*I*_NaK,max_ ∙{1+(K_mKp_/K_o_)^1.2^}^-1^ ∙ {1+(K_mNap_/Na_i_)^1.3^} ^-1^ ∙ {1+exp[-(*V_m_*- *E*_Na_+120)/30]}^-1^

**Ca^2+^- background current (*I*_bCa_)**

*I*_bCa_ = *C*_m_∙ *g*_bCa_ ∙(*V_m_* - *E*_CaL_)

**Na^+^-Ca^2+^ exchanger current (*I*_NCX_)**, based on original formulations from Dokos et al. (Dokos et al., 1996).

*I*_NCX_ = *C*_m_ ∙*k*_NCX_ ∙(*k*_21_ ∙ *x*_2_ - *k*_12_ ∙ *x*_1_) / (*x*_1_ + *x*_2_ + *x*_3_ + *x*_4_)

*d*_o_ =1+(*Ca*_o_/*K*_co_)∙{1+exp(*Q*_co_∙*V_m_*/*E*_T_)}+(*Na*_o_/*K*_1no_)∙{1+(*Na*_o_/*K*_2no_)∙(1+*Na*_o_/*K*_3no_)}

*k*_43_ = *Na*_i_/(*K*_3ni_ +*Na*_i_)

*k*_41_ = exp[-*Q*_n_∙*V_m_*/(2*E*_T_)]

*k*_34_ = *Na*_o_/(*K*_3no_+*Na*_o_)

*k*_21_ = (*Ca*_o_/*K*_co_)∙exp(*Q*_co_∙*V_m_*/*E*_T_) /*d*_o_

*k*_23_ = (*Na*_o_/*K*_1no_)∙(*Na*_o_/*K*_2no_)∙(1+*Na*_o_/*K*_3no_)∙ exp[-*Q*_n_∙*V_m_*/(2*E*_T_)]/*d*_o_

*k*_32_ = exp[*Q*_n_∙*V_m_*/(2*E*_T_)]

*x*_1_ = *k*_34_ ∙ *k*_41_ ∙(*k*_23_ + *k*_21_) + *k*_21_ ∙ *k*_32_ ∙(*k*_43_ + *k*_41_)

*d*_i_ = 1+(*Ca*_sub_/*K*_ci_)∙{1+exp(-*Q*_ci_∙*V_m_*/*E*_T_)+*Na*_i_/*K*_cni_}+(*Na*_i_/*K*_1ni_)∙{1+(*Na*_i_/*K*_2ni_)∙(1+*Na*_i_/*K*_3ni_)}

*k*_12_ =(*Ca*_sub_/*K*_ci_)∙exp(-*Q*_ci_∙*V_m_*/*E*_T_)/*d*_i_

*k*_14_ = (*Na*_i_/*K*_1ni_)∙(*Na*_i_/*K*_2ni_)∙(1 +*Na*_i_/*K*_3ni_)∙ exp[*Q*_n_∙*V_m_*/(2*E*_T_)]/*d*_i_

*x*_2_ = *k*_43_∙ *k*_32_ ∙(*k*_14_ + *k*_12_) + *k*_41_∙ *k*_12_ ∙*k*_34_ + *k*_32_)

*x*_3_ = *k*_43_ ∙*k*_14_ ∙(*k*_23_ + *k*_21_) + *k*_12_ ∙*k*_23_∙(*k*_43_ + *k*_41_)

*x*_4_ = *k*_34_ ∙*k*_23_ ∙(*k*_14_ + *k*_12_) + *k*_21_ ∙*k*_14_∙(*k*_34_+ *k*_32_)

**Acetylcholine-activated K^+^ current (*I*_KACh_),** adopted from Demir et al. 1999 (Demir et al., 1999) (Note ***I*_KACh_** = 0 when [ACh]=0)

*I*_KACh_ = *a* ⋅ *g*_KACh,max_ ⋅ (*V*_m_ - *E*_K_)

*beta*=0.001 · 12.32/(1+0.0042/[ACh]) (per ms)

*alfa*= 0.001 · 17⋅exp(0.0133⋅(*V_m_*+40)) (per ms)

*a*_∞_ = *beta* / (*alfa* + *beta*)

*τ*_a_ = 1/(*alfa* + *beta*) (in ms)

**FORMULATIONS: Ca^2+^ CYCLING**

**Ca^2+^ release flux (*j*_SRCarel_) from SR via RyRs**, based on original formulations of Stern et al. (Stern et al., 1999) and modified by Shannon et al. (Shannon et al., 2004)

*j*_SRCarel_ = *k*_s_∙*O*∙(*Ca*_jSR_ - *Ca*_sub_)

*k*_CaSR_ = *MaxSR*- (*MaxSR* - *MinSR*)/ (1 + (*EC*_50_SR_/*Ca*_jSR_)^HSR^)

*k*_oSRCa_ = *k*_oCa_/*k*_CaSR_

*k*_iSRCa_ = *k*_iCa_∙*k*_CaSR_

*dR/dt* = (*k*_im_∙*RI* - *k*_iSRCa_ ∙*Ca*_sub_∙*R*) - (*k*_oSRCa_∙*Ca*_sub_^2^∙*R* - *k*_om_∙*O*)

*dO/dt* =(*k*_oSRCa_∙*Ca*_sub_^2^ ∙*R* - *k*_om_∙*O*) - (*k*_iSRCa_∙*Ca*_sub_∙*O* - *k*_im_∙*I*)

*dI/dt* = (*k*_iSRCa_∙ *Ca*_sub_∙*O* - *k*_im_∙*I*) - (*k*_om_∙*I* - *k*_oSRCa_∙*Ca*_sub_^2^ ∙*RI*)

*dRI/dt* = (*k*_om_∙*I* - *k*_oSRCa_∙*Ca*_sub_^2^∙*RI*) - (*k*_im_∙*RI* - *k*_iSRCa_∙*Ca*_sub_∙*R*)

**Intracellular Ca^2+^ fluxes**

**Ca^2+^ diffusion flux** **(*j*_Ca_dif_)** from submembrane space to myoplasm:

*j*_Ca_dif_ = (*Ca*_sub_ - *Ca*_i_)/τ_difCa_

**The rate of Ca^2+^ uptake (pumping) (*j*_up_)** by the SR, based on formulations of SR Ca^2+^ pump function suggested by Luo and Rudy (Luo and Rudy, 1994).

*j*_up_ = *P*_up_ /(1 + *K*_up_/*Ca*_i_)

Autonomic modulation of Ca^2+^ uptake was described as reported previously (Maltsev and Lakatta, 2010): β-AR stimulation doubled *P*_up_ from *P*_up,basal_ of 0.012 mM/ms to 0.024 mM/s; by ChR stimulation inhibited *P*_up_, with fractional block (*b*_up_) as follows:

*P*_up_ = *P*_up,basal_ · (1 - *b*_up_)

*b_up_ = b_up,max_* ⋅ [ACh]/( *K_0.5,up_* + [ACh])

where *K_0.5,up_* = 90 nM is the [ACh] for half-maximal inhibition and *b_up,max_* = 0.7.

**Ca^2+^ flux between (network and junctional) SR compartments (*j*_tr_)**:

*j*_tr_ = (*Ca*_nSR_ – *Ca*_jSR_)/τ_tr_

**Ca^2+^ buffering**

*df*_TC_*/dt* = *k*_fTC_∙*Ca*_i_∙(1 -*f*_TC_) - *k*_bTC_ ∙ *f*_TC_

*df*_TMC_/*dt* = *k*_fTMC_ ∙*Ca*_i_ ∙(1- *f*_TMC_ - *f*_TMM_) - *k*_bTMC_ ∙ *f*_TMC_

*df*_TMM_/*dt* = *k*_fTMM_ ∙*Mg*_i_ ∙(1-*f*_TMC_ - *f*_TMM_)- *K*_bTMM_ ∙ *f*_TMM_

*df*_CMi_/*dt* =	*k*_fCM_ ∙*Ca*_i_ ∙(1- *f*_CMi_) - *k*_bCM_ ∙ *f*_CMi_

*df*_CMs_/*dt* = *k*_fCM_ ∙*Ca*_sub_∙(1 - *f*_CMs_) - *k*_bCM_ ∙ *f*_CMs_

*df*_CQ_/*dt* =	*k*_fCQ_ ∙*Ca*_jSR_∙(1- *f*_CQ_) - *k*_bCQ_ ∙ *f*_CQ_

**Dynamics of Ca^2+^ concentrations in cell compartments**

*dCa*_i_/*dt* =(*j*_Ca_dif_ ∙*V*_sub_ - *j*_up_∙ *V*_nSR_) /*V*_i_ - (*CM*_tot_∙*df*_CMi_*/dt* + *TC*_tot_∙*df*_TC_/*dt* + *TMC*_tot_∙*df*_TMC_*/dt*)

*dCa*_sub_*/dt* = *j*_SRCarel_ ∙*V*_jSR_/*V*_sub_ -(*I*_CaL_+*I*_CaT_+*I*_bCa_-2∙*I*_NCX_)/(2∙F∙*V*_sub_)-(*j*_Ca_dif_ + *CM*_tot_ ∙*df*_CMs_*/dt*)

*dCa*_jSR_/*dt* = *j*_tr_ - *j*_SRCarel_ - *CQ*_tot_ ∙ *df*_CQ_*/dt*

*dCa*_nSR_*/dt* = *j*_up_ - *j*_tr_ ∙*V*_jSR_/*V*_nSR_

**Initial values**

Online Table S5 summarizes all model variables (*y_1_*–*y_30_*) with their initial values.

**Model modification to generate AP firing interval (APFI) variability**

The original Maltsev-Lakatta model could not be directly used for APFI variability simulations because it is a system of 30 first-order differential equations (ODEs) that is deterministic and showing no APFI variability in limit cycle oscillatory regime of AP steady firing. Thus, we modified the model to generate variability of AP waveforms by supplementing total membrane current (I_tot_) with an additional current randomly fluctuating around its zero-mean value, known as perturbation current or I_per_ (as previously implemented by Henggui Zhang {Monfredi, 2014 #202}). Computer code implementation for I_per_ was as follows: I_per_ = I_per,max_* *ξ(t),* where I_per,max_ was the maximum amplitude of the perturbing current and ξ(t) is a random fluctuation term (‐1≤ξ(t)≤+1) that was generated by a random number generator within the program code. Random numbers were generated every 4 ms. I_per,max_ was tuned for the APFI variability in the model to match that measured experimentally under respective experimental conditions. We investigated two scenarios of noise generation when I_per_ was added to either I_tot_ or Ca release flux *j*_SRCarel_ (I_tot_ in pA was recalculated in terms of mM/ms) in three conditions (i) basal AP firing, (ii) ISO 100 nM, and (iii) CCh (0.1 μM) (Figures S8 and S9). Variability of 6 major currents was measured and analyzed: I_f_, I_NCX_, I_Kr_, I_CaL_, I_CaT_ and I_KACh_. We also measured variability of [Ca] under cell membrane. For all items we measured variability of their peak amplitudes and amplitudes at -40 mV during DD. An example of analysis for I_NCX_ is shown Figure S10 and all results are summarized in Tables S6 and S7.

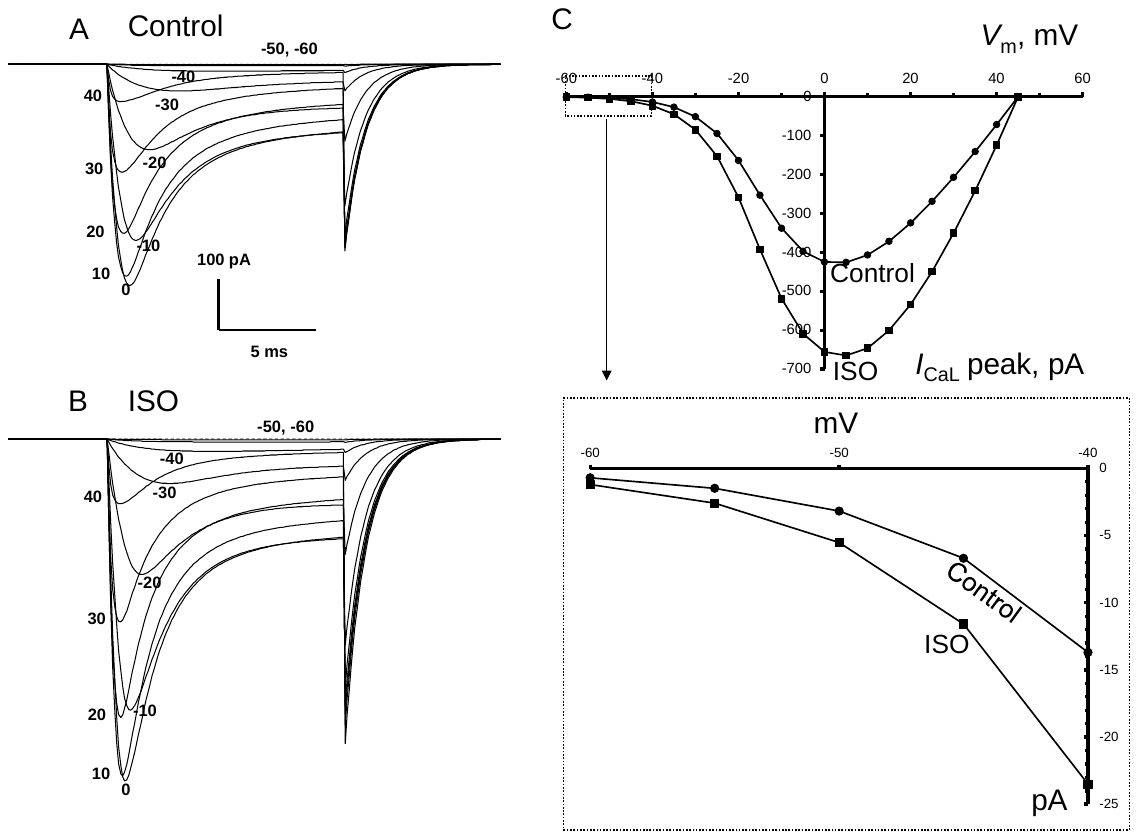

**Supplementary Figure S6**. Model simulations of β-AR stimulation effect on current-voltage relationship for *I*_CaL_. *I*_CaL_ traces were simulated by applying testing voltage pulses *V_m_* from a holding potential of -80 mV. A and B show simulated traces for *V_m_* from -60 mV to 40 mV with a 10 mV interval in the basal state (Control), in the presence of β-AR stimulation by ISO. The values of *V_m_* are shown by labels at the respective current peaks. D: The current voltage relationships (5 mV voltage step) for *I*_CaL_ peak. Inset illustrates presence of *I*_CaL_ current activation in the model within the voltage range of the diastolic depolarization from -60 mV to -40 mV. Cell electric capacitance is 32 pF

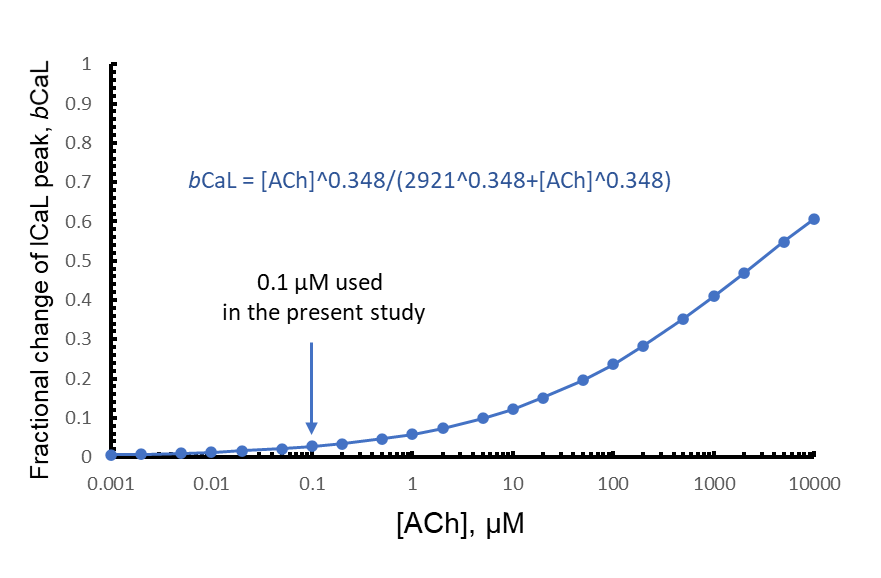

**Supplementary Figure S7**. Fractional block (*b*_CaL_) of *I*_CaL_ by ChR stimulation adopted from Zaza et al. model given in Figure 2 legend in (Zaza et al., 1996). The fractional block in the presence of 0.1 μM of CCh (simulated in the present study) is expected relatively small: *b*_CaL_(0.1) = 0.02728, i.e. <3%.

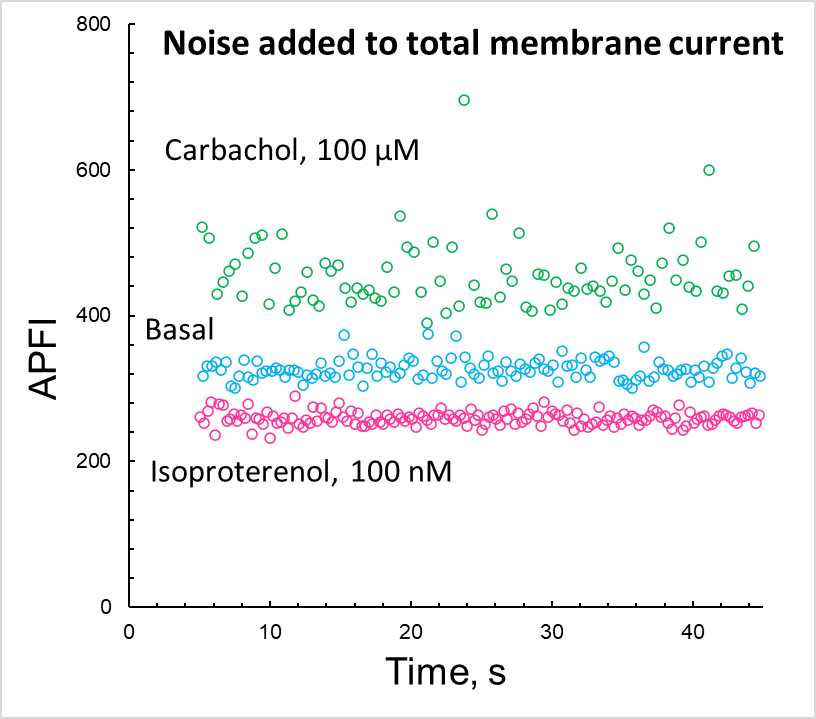

**Supplementary Figure S8.** Intervalogram of AP firing simulated by coupled-clock Maltsev-Lakatta model with noise (I_per_) added to total current (I_tot_). I_per,max_ (pA) was tuned for the APFI variability in the model to match that measured experimentally: 15 pA, 10, and 5 pA, in ISO, Basal, and CCh, respectively.

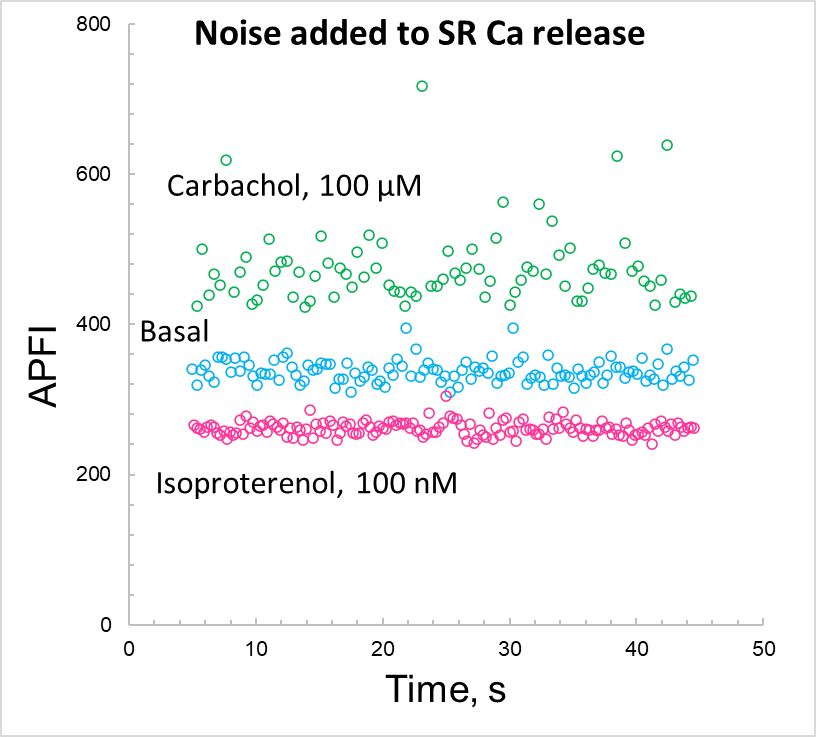

**Supplementary Figure S9**. Intervalogram of AP firing simulated by coupled-clock Maltsev-Lakatta model with noise (I_per_) added to SR Ca release flux. I_per,max_ (pA) was tuned for the APFI variability in the model to match that measured experimentally: 33, 28, and 15 pA, in ISO, Basal, and CCh, respectively.

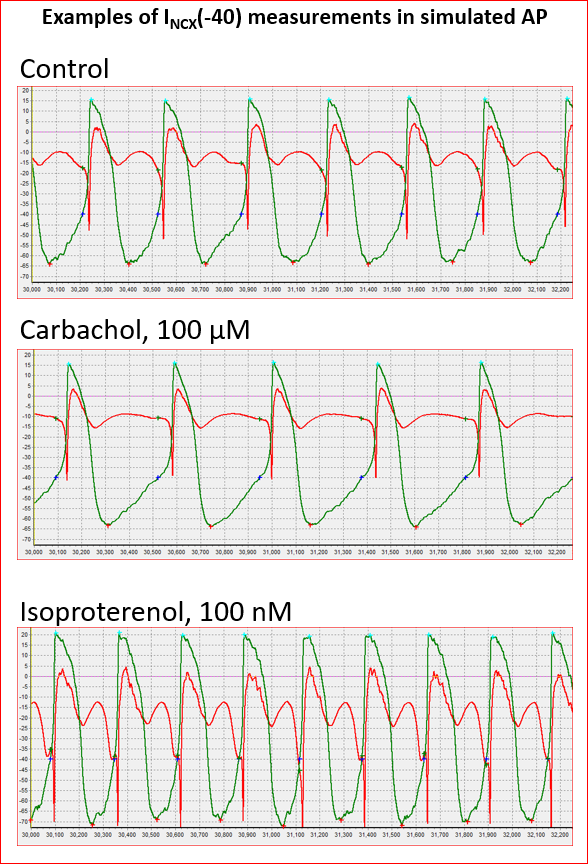

Time, ms

**Supplementary Figure S10**. An example of simulations and analysis of AP firing (green traces) and I_NCX_ (red traces) with noise current (I_per_) added to total current I_tot_ to match respective CV of APFI measured experimentally. Crosses: red is maximum diastolic potentials; aqua is AP overshoot, blue is -40 mV, green I_NCX_ value at -40 mV.

**Supplementary Table S4.**

Parameters of steady-state activation and inactivation for *I*_CaL_ in the present model compared to previous rabbit SANC models (see main text Methods for details). * Steady state inactivation parameters were set in our model to the respective values measured in our laboratory in an experimental study of isolated rabbit SANC by Vinogradova et al. (Vinogradova et al., 2000).

| Model | Steady-state activation curve,  *d*_L,∞_ | | Steady-state inactivation curve,  *f*_L,∞_ | |
| --- | --- | --- | --- | --- |
|  | Midpoint,  *V*_½,d_ (mV) | Slope factor  *K*_d_ (mV) | Midpoint,  *V*_½,f_ (mV) | Slope factor  *K*_f_ (mV) |
| Wilders et al. (Wilders et al., 1991) | -6.6 | 6.6 | -25 | 6 |
| Demir et al. (Demir et al., 1994) | -14.1 | 6 | -25 | 5 |
| Dokos et al. (Dokos et al., 1996) | -6.6 | 6.6 | -25 | 6 |
| Zhang et al. (Zhang et al., 2000) | -23.1 | 6 | -45 | 5 |
| Kurata et al. (Kurata et al., 2002) | -14.1 | 6 | -30 | 5 |
| **Present model** | **-13.5** | **6** | **-35*** | **7.3*** |

**Supplementary Table S5.** Model variables: description and initial values. Our SAN cell model is described as a system of 30 first order differential equations (variables *y_1_ - y_30_*).

| # | Variable | Description | Initial value |
| --- | --- | --- | --- |
| ***Ca^2+^ cycling*** | | | |
| *y_1_* | *Ca_i_* | [Ca^2+^] in myoplasm, mM | 0.0001 |
| *y_2_* | *Ca_sub_* | [Ca^2+^] in submembrane space, mM | 0.000223 |
| *y_3_* | *Ca*_jSR_ | [Ca^2+^] in the junctional SR (jSR), mM | 0.029 |
| *y_4_* | *Ca*_nSR_ | [Ca^2+^] in the network SR (nSR), mM | 1.35 |
| *y_5_* | *f*_TC_ | Fractional occupancy of the troponin-Ca^2+^ site by Ca^2+^ in myoplasm | 0.02 |
| *y_6_* | *f*_TMC_ | Fractional occupancy of the troponin-Mg^2+^ site by Ca^2^ in myoplasm | 0.22 |
| *y_7_* | *f*_TMM_ | Fractional occupancy of the troponin-Mg^2+^ site by Mg^2+^ in myoplasm | 0.69 |
| *y_8_* | *f*_CMi_ | Fractional occupancy of calmodulin by Ca^2+^ in myoplasm | 0.042 |
| *y_9_* | *f*_CMs_ | Fractional occupancy of calmodulin by Ca^2+^ in submembrane space | 0.089 |
| *y_10_* | *f*_CQ_ | Fractional occupancy of calsequestrin by Ca^2+^ in junctional SR | 0.032 |
| *y_11_* | *R* | RyR reactivated (closed) state | 0.7499955 |
| *y_12_* | *O* | RyR open state | 3.4·10^-6^ |
| *y_13_* | *I* | RyR inactivated state | 1.1·10^-6^ |
| *y_14_* | *RI* | RyR RI state | 0.25 |
| ***Electrophysiology*** | | | |
| *y_15_* | *V_m_* | Membrane potential, mV | -65 |
| *y_16_* | *d*_L_ | *I*_CaL_ activation | 0 |
| *y_17_* | *f*_L_ | *I*_CaL_ voltage-dependent inactivation | 1 |
| *y_18_* | *f*_Ca_ | *I*_CaL_ Ca^2+^ dependent inactivation | 1 |
| *y_19_* | *p*_aF_ | *I*_Kr_ fast activation | 0 |
| *y_20_* | *p*_aS_ | *I*_Kr_ slow activation | 0 |
| *y_21_* | *p*_i_ | *I*_Kr_ inactivation | 1 |
| *y_22_* | *n* | *I*_Ks_ activation | 0 |
| *y_23_* | *y* | *I*_f_ activation | 1 |
| *y_24_* | *d*_T_ | *I*_CaT_ activation | 0 |
| *y_25_* | *f*_T_ | *I*_CaT_ inactivation | 1 |
| *y_26_* | *q* | *I*_to_ inactivation | 1 |
| *y_27_* | *r* | *I*_to_ and *I*_sus_ activation | 0 |
| *y_28_* | *q_a_* | *I*_st_ activation | 0 |
| *y_29_* | *q_i_* | *I*_st_ inactivation | 1 |
| *y_30_* | *a* | *I*_KACh_ activation | 1 |

**Supplementary Table S6**. Results of variability analysis of major ion currents I_f_, I_Kr_, I_CaL_, I_CaT_, I_KACh_, and submembrane Ca simulated by a coupled-clock model **with noise added to total membrane current (I_tot_)** to match APFI variability observed experimentally. The results are presented for control, β-adrenergic receptor stimulation (ISO, 100 nM), and cholinergic receptor stimulation (CCh 0.1 μM) separately for peak amplitudes and amplitudes at -40 mV at the middle of diastolic depolarization. SD, standard deviation; CV, coefficient of variation (Mean/SD in %). The simulations were 40 s long with the total # of cycles indicated in parentheses. I_per,max_ (pA) was tuned for the APFI variability in the model to match that measured experimentally: 15 pA, 10, and 5 pA, in ISO, Basal, and CCh, respectively.

|  | ISO (154 cycles) | | | Basal (122 cycles) | | | CCh (87 cycles) | | |
| --- | --- | --- | --- | --- | --- | --- | --- | --- | --- |
|  | Mean | SD | CV, % | Mean | SD | CV, % | Mean | SD | CV, % |
| APFI (ms) | 259.25 | 9.1321 | 3.5225 | 326.45 | 13.90 | 4.257 | 455.78 | 45.562 | 9.997 |
| **The peak amplitudes of simulated ion currents** | | | | | | | | | |
| I_NCX_ (pA) | -74.32 | 3.651 | 4.9128 | -48.52 | 2.498 | 5.148 | -39.11 | 1.893 | 4.840 |
| I_f_ (pA) | -3.667 | 0.2350 | 6.4082 | -2.163 | 0.1937 | 8.953 | -1.554 | 0.0780 | 5.018 |
| I_Kr_ (pA) | 90.545 | 0.6646 | 0.734 | 56.76 | 0.4128 | 0.727 | 54.55 | 0.3655 | 0.670 |
| I_CaL_ (pA) | -313.5 | 4.9296 | 1.572 | -212.2 | 4.416 | 2.081 | -205.7 | 6.212 | 3.019 |
| I_CaT_ (pA) | -18.69 | 5.6899 | 30.447 | -3.841 | 0.8659 | 22.55 | -2.109 | 0.1317 | 6.248 |
| Ca_sub_ (μM) | 2.5616 | 0.0986 | 3.8486 | 1.818 | 0.0647 | 3.557 | 1.566 | 0.0513 | 3.276 |
| I_KACh_ (pA) |  |  |  |  |  |  | 6.809 | 0.1323 | 1.943 |
| **The values of simulated currents measured at a membrane potential of -40mV** | | | | | | | | | |
| I_NCX_ (pA) | -40.53 | 3.803 | 9.383 | -17.81 | 1.0768 | 6.047 | -10.28 | 0.7026 | 6.837 |
| I_f_ (pA) | -1.883 | 0.0857 | 4.553 | -1.291 | 0.0784 | 6.068 | -0.938 | 0.0284 | 3.029 |
| I_Kr_ (pA) | 32.95 | 0.8466 | 2.5694 | 19.66 | 0.5984 | 3.043 | 16.208 | 0.3041 | 1.876 |
| I_CaL_ (pA) | -11.66 | 0.4961 | 4.2562 | -8.688 | 0.2449 | 2.819 | -9.081 | 0.1838 | 2.0245 |
| I_CaT_ (pA) | -10.32 | 1.5963 | 15.461 | -3.841 | 0.8659 | 22.55 | -1.334 | 0.1878 | 14.075 |
| [Ca^2+^] under membrane(μM) | 0.5213 | 0.3625 | 6.954 | 0.3134 | 0.00932 | 2.977 | 0.2491 | 0.0059 | 2.371 |
| I_KACh_ (pA) |  |  |  |  |  |  | 3.8294 | 0.0615 | 1.6061 |

**Supplementary Table S7**. Results of variability analysis of major ion currents I_f_, I_Kr_, I_CaL_, I_CaT_, I_KACh_, and submembrane Ca simulated by a coupled-clock model **with noise added to Ca release flux** to match APFI variability observed experimentally. The results are presented for control, β-adrenergic receptor stimulation (ISO, 100 nM), and cholinergic receptor stimulation (CCh, 0.1 μM) separately for peak amplitudes and amplitudes at -40 mV, i.e. at the middle of diastolic depolarization. SD, standard deviation; CV, coefficient of variation (Mean/SD in %). The simulations were 40 s long with the total # of cycles indicated in parentheses. I_per,max_ (pA) was tuned for the APFI variability in the model to match that measured experimentally: 33, 28, and 15 pA, in ISO, Basal, and CCh, respectively.

|  | ISO (152 cycles) | | | Basal (118 cycles) | | | CCh (83 cycles) | | |
| --- | --- | --- | --- | --- | --- | --- | --- | --- | --- |
|  | Mean | SD | CV, % | Mean | SD | CV, % | Mean | SD | CV, % |
| APFI (ms) | 261.44 | 9.321 | 3.565 | 337.43 | 14.61 | 4.331 | 474.59 | 50.42 | 10.62 |
| **The peak amplitudes of simulated ion currents** | | | | | | | | | |
| I_NCX_ (pA) | -74.33 | 3.098 | 4.167 | -47.95 | 2.879 | 6.004 | -38.42 | 2.142 | 5.576 |
| I_f_ (pA) | -3.710 | 0.2739 | 7.382 | -2.143 | 0.1536 | 7.166 | -1.542 | 0.0704 | 4.564 |
| I_Kr_ (pA) | 90.50 | 0.2520 | 0.2784 | 56.70 | 0.2053 | 0.362 | 54.39 | 0.3704 | 0.681 |
| I_CaL_ (pA) | -313.6 | 4.169 | 1.329 | -211.9 | 3.212 | 1.516 | -205.0 | 6.966 | 3.398 |
| I_CaT_ (pA) | -18.98 | 8.052 | 42.43 | -3.312 | 0.7169 | 21.64 | -2.015 | 0.1379 | 6.841 |
| Ca_sub_ (μM) | 2.551 | 0.079 | 3.096 | 1.815 | 0.0700 | 3.855 | 1.556 | 0.0592 | 3.802 |
| I_KACh_ (pA) |  |  |  |  |  |  | 6.779 | 0.1272 | 1.875 |
| **The values of simulated currents measured at a membrane potential of -40mV** | | | | | | | | | |
| I_NCX_ (pA) | -42.80 | 8.284 | 19.36 | -18.49 | 4.359 | 23.57 | -11.72 | 1.887 | 16.10 |
| I_f_ (pA) | -1.919 | 0.1128 | 5.881 | -1.321 | 0.0591 | 4.475 | -0.950 | 0.0357 | 3.755 |
| I_Kr_ (pA) | 32.78 | 0.5673 | 1.731 | 19.49 | 0.3704 | 1.900 | 16.04 | 0.3277 | 2.043 |
| I_CaL_ (pA) | -11.67 | 0.7497 | 6.422 | -8.797 | 0.2594 | 2.949 | -9.072 | 0.1961 | 2.162 |
| I_CaT_ (pA) | -10.36 | 2.447 | 24.14 | -2.951 | 0.7586 | 25.71 | -1.290 | 0.1494 | 11.58 |
| [Ca^2+^] under membrane(μM) | 0.5442 | 0.0818 | 15.02 | 0.3198 | 0.03796 | 11.87 | 0.2613 | 0.0160 | 6.109 |
| I_KACh_ (pA) |  |  |  |  |  |  | 3.806 | 0.0600 | 1.577 |
